# Quantitative analyses of single mitochondrial structure-function heterogeneity uncovers stemness specification by redox-tuned small-mitochondrial-networks

**DOI:** 10.1101/2024.12.26.630414

**Authors:** Mayank Saini, Swati Agarwala, Biratal Wagle, Bharti Golchha, Brian Spurlock, Pranit Sinha, Partha Pratim Chakrabarti, Diya Prasad, Danitra Parker, Akshara Kulkarni, Kasturi Mitra

## Abstract

Lack of quantitative understanding of the marked heterogeneity of multifaceted mitochondria poses challenges in unraveling their translatable role. Here, we hierarchically untangled multilevel heterogeneity of mitochondrial networks and non-networks by simultaneously analysing their structure-function within single components and their units, in cells and tissues. Such mito-SinComp quantitative analyses revealed that redox levels of mitochondrial networks and their dramatic intra-network heterogeneity can be predicted by structural features through a translatable machine-learning approach. Mito-SinComp identified and quantitatively characterized a redox-tuned subpopulation of Small-Mitochondrial-Networks(SMNs) that specifies stemness. These SMNs are generated by severing of nodes of oxidized Hyperfused-Mitochondrial-Networks(HMNs), are ten-folds smaller with specific network complexity and elevated mt-DNA nucleoid abundance. Thus, HMN to SMN conversion supports elevated expression of mtDNA genes in establishing MT-ND1(redox)-KRT15(stemness) transcriptomic interaction, as revealed by coupled scRNA-seq. This stemness priming of non-transformed cells is sustained after neoplastic transformation and is detected in patient transcriptome, demonstrating translational relevance of SMNs.

## Introduction

Mitochondria, as dynamic organelles, are gatekeepers of multiple cellular homeostatic processes and form signalling hubs^1,2^. Mitochondria can be heterogenous between cells, tissues, and also individuals, while a cell can have few to hundreds of mitochondria of heterogenous structures including their elaborate physical networks. Reorganization of mitochondrial structure by cellular processes is linked to various disease etiologies^3–7^.

Mitochondrial structure-function is primarily modulated by the counteracting processes of mitochondrial fission and fusion, and extensive molecular characterization of such processes is an active research area ^6,8^. Network forming ability of mitochondria regulates fundamental mitochondrial processes and may have imparted evolutionary advantage ^9–13^. However, various fundamental questions remain about how non-networked and networked mitochondrial entities interact to crosstalk with mitochondrial function and thus impact cell physiology.

Lack of quantitative knowledge of mitochondrial network heterogeneity poses challenges in developing generalized understanding of the function of mitochondrial networks, shrouding the field with controversies. Additionally, the natural heterogeneity of mitochondrial structure-function along a wide spectrum is not reflected accurately by genetic alteration of the molecular regulators. Therefore, genetic approaches sometimes lead to confusing findings preventing building knowledge consensus of the exact role of mitochondria. Moreover, genetic experiments may obscure the distinction between primary and secondary impacts due to remodelling ability of mitochondria. Therefore, there is a serious necessity of obtaining direct quantitative understanding of the heterogeneity of mitochondrial structure-function across their natural spectrum towards elucidating the interplay of mitochondria with cellular process. Various laboratories, including our own, have attempted to quantitatively define mitochondrial structure using microscopy^14–22^. We reported that studying mitochondrial structure-function in single cells characterizes mitochondria primed stemness states, similar to single cell transcriptomics^21,23,24^.

Contradictory reports about the mitochondrial structure and number in stem cells clouds our understanding of the impact of mitochondria on stem cell physiology ^3,25–27^. Lineage specific adult stem cells undergo neoplastic transformation into tumor initiating cells, and some are particularly vulnerable to mitochondrial inhibition ^28–30^. Stemness state (characterized by self-renewal) can be sustained by repression of mitochondrial fission ^3,21,23,24,31,32^, while stem cells are generally considered to have rudimentary fragmented mitochondrial structures ^33,34^.

However, our quantitative approach revealed repression of mitochondrial fission as well as its regulated activation are required to prime a stem cell state to initiate and sustain neoplasticity ^23^. Activation of mitochondrial fission is essential for oncogene induced neoplastic transformation ^35,36^. A recent report proposed that the mitochondrial fission and fusion cycles modulate oxidative and reductive biosynthesis in same mitochondrial populations ^5^, highlighting mitochondria fission and fusion act in concert in regulating distinct aspects of a given cellular process.

We and others previously identified the hyperfused mitochondria in certain cellular processes, which is primarily regulated by repression of mitochondrial fission ^22,23,37,38^. Here, we hierarchically untangled inter- and intra-cellular as well as intra-network mitochondrial heterogeneity by directly quantifying mitochondrial structure-function of single mitochondrial components. This mito-SinComp analyses revealed novel concepts in mitochondrial redox physiology and mtDNA functional organization and their mechanistic role in stem cell physiology. We identified and quantitatively characterized a subpopulation of small mitochondrial networks (SMNs), which are derived from the hyperfused mitochondrial networks (HMNs) to specify stemness. Using coupled single cell transcriptomics we identified and validated key molecules underlying the SMN driven stemness specification, which is also detectable in patients. Our data demonstrate that unprecedented resolution of mitochondrial structure-function heterogeneity, enabled by mito-SinComp, allows fundamental discovery and its translational application.

## Results

### Modular design of mito-SinComp analyses of single mitochondrial components for hierarchical untangling of mitochondrial structure(S)-function(F) heterogeneity

Our single cell microscopy based approach of mitochondrial heterogeneity analyses has revealed novel findings about quantitatively defined mitochondrial structural states in guiding cell states ^23,39^. Here, we unrestricted such analyses to study mitochondrial structure-function heterogeneity in a ‘hierarchical manner’ for ‘any’ given mitochondrial function in ‘any’ given cell physiology using a singular quantitative approach. This is a fluorescence pixel level simultaneous quantitative analyses of structure-function of single mitochondrial components and its units for live or fixed cells / tissues, namely mito-SinComp. Such analyses can untangle mitochondrial structure-function heterogeneity between and within individual subjects→ tissues→ cells→ mitochondrial networks or non-networks→ their types and subtypes (based on their features) →elements forming the networks→zones forming the elements **(Fig. 1).**

**Figure. 1:**
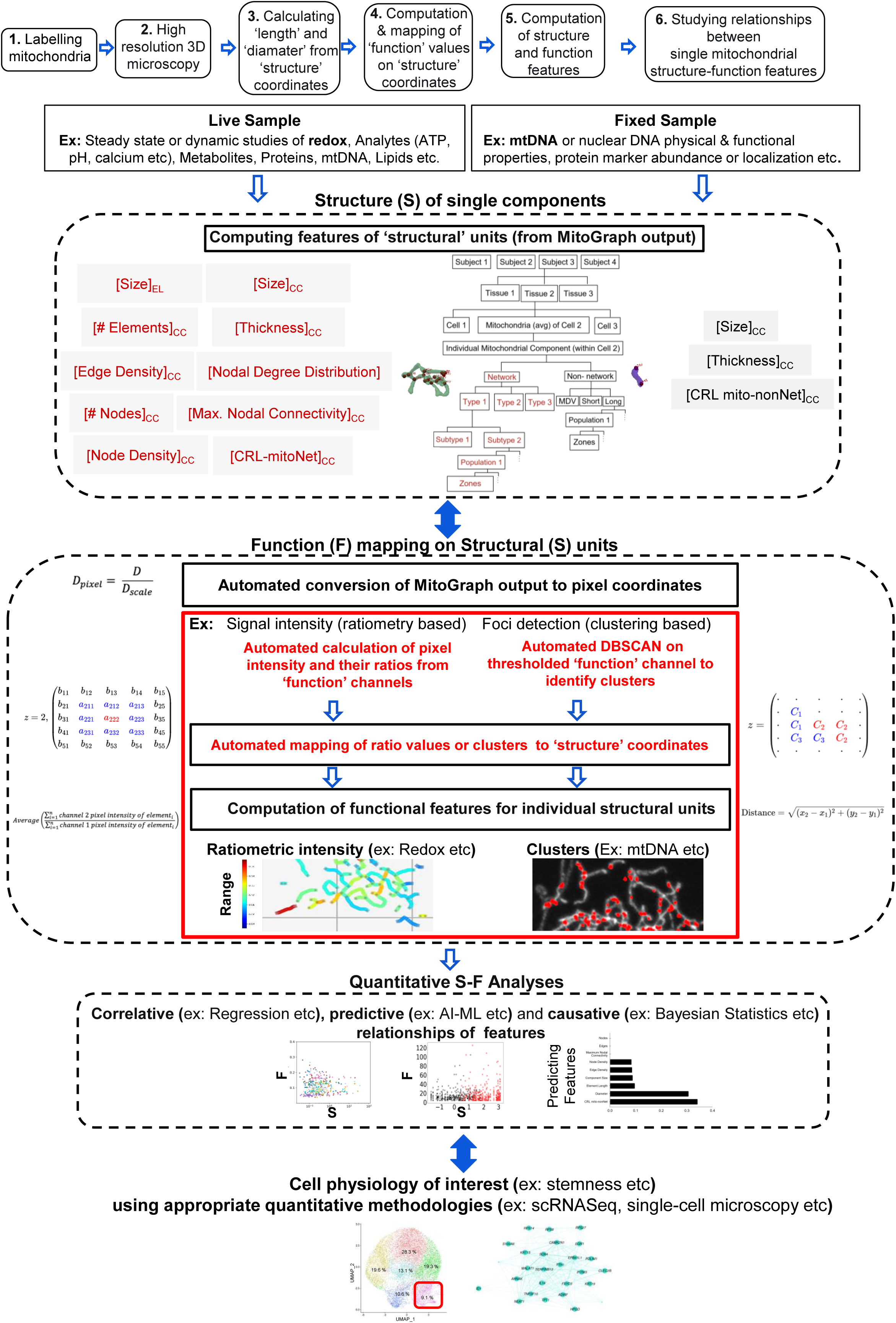
Modular design of quantitative mito-SinComp approach. Mitochondrial heterogeneity can be untangled at hierarchical levels following 6 steps in live and fixed cells or tissues. Multiple structural features (S) are extracted and the function (F) is computed and mapped on the function. S-F analyses of single mitochondrial components can be performed with various statistical methods. Standalone modular use of mito-SinComp to elucidate mitochondrial physiology is demonstrated through examples of redox function and functional mtDNA organization. Combinatorial use of mito-SinComp to study mitochondrial heterogeneity in the cell physiology is demonstrated using scRNA-seq modality to study stemness.

The approach involves the following steps on the total mitochondrial population in live or fixed cells or tissues, within or beyond cellular boundaries (details in Methods): **1)** combinatorial labelling of mitochondria with a structure probe and one or more compatible function probes (chemical, immuno- or genetic); **2)** 3D fluorescence confocal microscopy to image the total population of mitochondria within a cell or tissue allowing wider applicability; **3)** application of MitoGraph software to obtain 3D ‘structure’ coordinates of individual mitochondrial components; **4)** automated mapping of any mitochondrial ‘function’ values on structure coordinates of all resolvable units for hierarchical computation of function; **5)** automated computation of length and diameter from pixel coordinates to compute the structure (S) and function (F) features for individual mitochondrial components (**Suppl Table 1**); **6)** studying bivariate or multivariate S-F or S-S or F-F relationships in associative, predictive or causal manner.

Modularity of mito-SinComp design allows use of any validated single or palate of mitochondrial functional probes for any mitochondrial sub-compartment, which are compatible with the structural probe of choice. Its scalability allows exploration with high-content, multiplexed, time-resolved or super-resolution microscopy, with appropriate modifications. Mito-SinComp performed in correlative or causative manner with other experimental modalities allows elucidation of significance of mitochondrial heterogeneity in any cell physiology. Mito-SinComp allows creation of library or map of mitochondrial S-F quantitative features across cells / tissues / organisms for extensive and intensive exploratory analyses, theoretical modelling with the mitochondrial S-F features and their relationships, and also deep mechanistic understanding of the mitochondrial properties and their heterogeneity in any given cell physiology. Although the primary goal of mito-SinComp does not include investigation of molecular mechanism, it can also be used in elucidation of the impact of genetic mutations or direct studies of mitochondrial regulators. The simplicity of the mito-SinComp approach allows fundamental and translational studies on drug, nutrient, macromolecule or genetic screening.

In the subsequent sections, we use mito-SinComp for hierarchical untangling of mitochondrial structure-function heterogeneity towards unraveling mechanistic role of mitochondria in specification of stemness state. We used 3 previously demonstrated stemness supporting conditions and tested in the situation of acute carcinogen exposure. We elucidate mitochondrial redox function in live cells and functional mtDNA organization within mitochondrial networks in fixed cells, and use quantitative mito-SinComp analyses in conjunction with quantitative single cell transcriptomics. The mitochondrial components (CC) having >1 mitochondrial edges (i.e. elements) connected by >2 nodes and with >2 terminal nodes are classified as ‘networked’, whereas mitochondrial components (CC) with 1 edge and 2 terminal nodes are classified as ‘non-networked’; the rare looped networked structures with one node and one edge are exceptions. The features with non-normal distributions are log-normalized and appropriate outlier filtering is included, as necessary. The numerical data from the automated analyses is collected in a comprehensive data file for downstream analyses, including statistical tests with multiple hypothesis testing adjustments. The physical properties of the networked mitochondria is studied using 10 structure features, while the non-networked mitochondria present fewer features, while the number of functional features may vary **(Fig. 1)**. Heterogeneity in the feature distribution is assessed by median and coefficient of variation while other statistical parameters of choice can be used.

### Extraction of quantitative S-F features of single mitochondrial components to study mitochondrial redox function in live cells

We used the HaCaT normal human keratinocyte model to study specification of stemness, where we demonstrated mitochondria priming of stemness supports neoplasticity driven by an oxidative carcinogen ^23^. After validating the mito-SinComp computed total component length of a cell with that of the MitoGraph output **(Fig. S1A)**, we tested the comprehensive mito-SinComp analytical script against our previous benchmark^23^. Mito-SinComp successfully identified the HaCaT populations harbouring the ‘intermediately-fused’ (**INT**) or the ‘hyperfused’ (**HF**) mitochondria in a pooled population where mitochondrial structure spectrum is widened with different levels of knockdown of the mitochondrial fission protein, Drp1, (**Fig. S1B**). Moreover, the 3D Graphs of the mitochondrial structural coordinates and the computed structural features for the identified networks and non-networks recapitulate the inter-and intra-cellular mitochondrial heterogeneity of the representative Cell 1-INT and Cell 2-HF (**Fig. 2A**, **Suppl. Table 2)**. Final validation of the mito-SinComp structural analyses comes from the successful detection of higher abundance of longer networked mitochondria (i.e. hyperfused) in the HF-Cells group **(Fig. 2B, left, arrow)**. The validated mito-SinComp structural analyses revealed the following quantitative comparisons between the INT-Cells and HF-Cells groups (**Fig. S1C-F**): **a)** comparable abundance of networks and non-networks per cell. **b)** networks are ∼10-100 times larger than the non-networks; **c)** average length of networks in individual cells is at least 2 folds less in the INT-Cells group in comparison to the HF-Cells group.

**Figure. 2:**
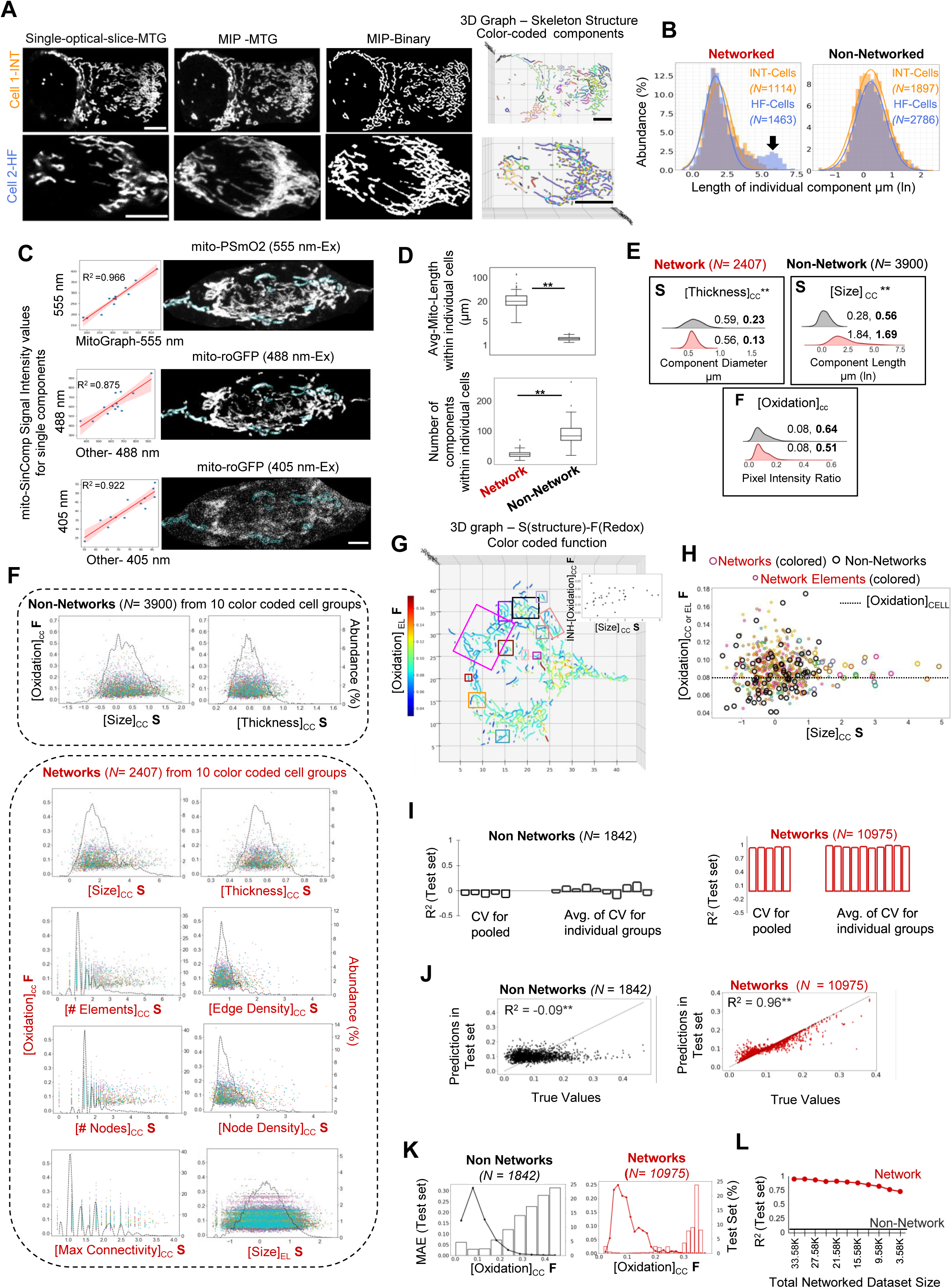
**(A)**. Labelled views of representative MTG stained single HaCaT cells. **(B)**. Histogram distribution of lengths of individual mitochondrial Networks and non-Networks (ignoring cell identity) for the colour-coded cell groups; arrow marks the hyperfused networks. **(C)**. Bivariate scatter plot of signal intensity values obtained with automated mito-SinComp and manually from ImageJ (left) from 13 mitochondrial components outlined in the representative cell expressing mito-PSMO2 and mito-roGFP (right). **(D)**. Box plot of average mitochondrial length and total number of mitochondrial Networks and non-Networks within individual cells in a pooled population of 10 experimental groups. **(E)**. Ridgeline plots comparing distributions of named (S) and (F) features between Networked and non-Networked mitochondria from (D); Median and Coefficient of Variation (in bold) are indicated for each feature. **(F)**. Bivariate S-F scatter plots of [Oxidation]_CC_ and individual structural features with their frequency distributions (dotted line) for Networked and non-Networked mitochondria from (D). **(G)**. Representative 3D S-F Graph of an HaCaT cell harboring mitochondrial components (outlined) in the median range of features; inset depicts bivariate S-F scatter plot of network [Size]_CC_ and INH-[Oxidation]_CC_ plot of each component. **(H)**. Bivariate S-F scatter plot of [Oxidation]_CC or EL_ and [Size]_CC_ for Networks and non-Networks for the cell in (G). **(I).** Bar plot of R^2^ values from ML based prediction of [Oxidation]_CC_ on the test set of the Networked and non-Networks from (D). **(J)**. Scatter plot of predicted [Oxidation] _CC_ values and True values from (I). **(K)**. Plot of [Oxidation]_CC_ prediction error (MAE) on test set (bar) with distribution of [Oxidation]_CC_ (line) for Networks and non-Networks. **(L)**. Plot of R^2^ values from ML based prediction of [Oxidation]_CC_ with reduction of total data set size for the Networks to the level of the non-Networks. **signifies Holmes-Bonferroni corrected p-value < 0.05; Scale bar: 5 μm; N = sample size.

Given stem cells are particularly dependent on redox tuning ^2,40^, we chose to study mitochondrial redox function that is bidirectionally linked with mitochondrial structure ^41^. We extracted the mito-SinComp structure and redox features from cells expressing our validated combination of genetically encoded mitochondrial structural probe mito-PSmO2 and general redox probe for mitochondrial matrix, mito-roGFP ^21^ **(Fig. 2C, right)**. First, mito-SinComp computes the [Oxidation]_Px_ values in the individual pixels and maps them to the mitochondrial coordinates identified in the binary image (**Fig. 1**, details in Methods). Thereafter, it compute the [Oxidation] values at different levels of structural hierarchically as follow: average of [Oxidation]_Px_ within an element yields [Oxidation]_EL_---average of [Oxidation]_EL_ of a component (network or non-network) yields [Oxidation]_CC_---average of [Oxidation]_CC_ within a cell yields the [Oxidation]_CELL_. Visually distinct mitochondrial components were chosen to validate the automated mito-SinComp intensity measurements by their strong linear correlation with values manually obtained with standard image analyses software **(Fig. 2C, left).** Weaker correlation of [Oxidation]_CELL_ values further indicates heterogeneity of intracellular mitochondrial signal (**Fig. S1G)**. Nonetheless, the refined mito-SinComp [Oxidation]_CELL_ values reveal statistically significant differences that are not the evident in manual analyses **(Fig. S1H).**

We demonstrated proof of principle of mito-SinComp S(network)-F(oxidation) analyses on a physiologically relevant heterogenous data set generated using multiple doses of an oxidative carcinogen, TCDD, to create a wider spectrum of [Oxidation] within non-transformed and transformed cells (control and carcinogen exposed). Mito-SinComp analyses of this pooled cell population of 106 cells from 6 experimental and 4 control groups, revealed that the average length of the networks in each cell ∼20 fold higher than that of the non-networks, and the average number of networks in each cell is ∼4 folds less than those non-networks **(Fig. 2D)**. Comparison of mito-SinComp S-F features shows that the networks are heterogenous in their [Size]_CC_ and non-networks are heterogenous in [Thickness]_CC_ and [Oxidation]_CC_ **(Fig. 2E)**. Individual group analyses reveal bimodal [Oxidation]_CC_ distribution in some and significant differences in [Oxidation] _CC_ between networks and non-networks in others **(Fig. S1I)**.

Thus, we developed and validated mito-SinComp analyses and extracted the mito-SinComp quantitative S(network) and F(redox) features from networks and non-networks of live cells. Using mito-SinComp on genetically or physiologically manipulated cell populations, we demonstrated the proof of principal of the approach in untangling of mitochondrial structure-function between networks and non-networks.

### Mitochondrial redox function and its heterogeneity can be predicted from mitochondrial network structure using mito-SinComp S-F features in machine learning algorithm

Of various possible analyses, we investigated the quantitative relationships between the single mitochondrial S-F features in steady state in the heterogenous pooled HaCaT populations as well as within individual cells. No obvious linear relationship is detected in the bivariate S-F analyses between [Oxidation]_CC_ and individual structural features of networked and non-networked mitochondrial components in the pooled data set (**Fig. 2F**). Notably, a complete range of [Oxidation]_CC_ and its intra-network heterogeneity is observed in the mitochondrial components in the median range of distribution of each structural feature **(Fig. 2F, Fig. S2A).** For analyses of intra-cellular heterogeneity, we identified a common cell that harbours mitochondrial components with S-F features in their respective median bins **(Fig. 2G)**. These mitochondrial components of this cell exhibit a variety of complex organization of mitochondrial elements within connected networks **(Fig. S2B).** Notably, the 3D S(structure)-F(redox) map of the cell reveals dramatic intra-network heterogeneity of mitochondrial redox levels between their constituent elements, quantified as INH-[Oxidation]_CC_ (noted in networks outlined in **Fig. 2G**, and quantified in inset, formula in Suppl Table 1). Interestingly, an S-F plot of [Size]_CC_ and [Oxidation]_CC_ and [Oxidation]_EL_ of their constituent elements reveal that the range of [Oxidation]_EL_ values are similar between networks and non-networks; average of these values yield [Oxidation]_CC_ of the respective network, which when averaged with the non-network yields [Oxidation]_CELL_ **(Fig. 2H).** Such intra-network redox heterogeneity may arise from maintenance of mitochondrial matrix discontinuity at the nodes, or by other undiscovered mechanisms.

Next, we investigated if the [Oxidation]_CC_ and INH-[Oxidation]_CC_ can be predicted from multiparametric analyses of their structural features. We used the Machine Learning (ML) method of Random Forest, to perform regression task towards predicting [Oxidation] levels considering the mitochondrial elements as the lowest structural unit; the high [Oxidation]_CC_ values consisting of 0.016 % of the total data set were filtered out as outliers. The training was performed with hyperparameter tuning in 70% of the data and tested in the remaining 30% of the data, with 5-fold cross validation (see Methods). The prediction was evaluated using R^2^ metric and prediction error was measured using Mean Absolute Error (MAE) metric.

Interestingly, [Oxidation]_CC_ levels can be robustly predicted for the mitochondrial networks (cross validated R^2^ of test scores = 0.957) but not for the mitochondrial non-networks, both using the pooled or individual experimental groups **(Fig. 2I,J)**. The INH-[Oxidation]_CC_ for the networks is also predictable to the same degree (cross validated R^2^ of test scores = 0.937). Comparison of MAE and [Oxidation]_CC_ distribution in the test set revealed that the prediction error increases with reduction in data abundance, with overall higher error for the non-networks **(Fig.2K)**. Given, the non-network data set is ∼6 fold smaller than that of the networks (due to a singular element), we tested the performance of the model after randomized reduction of the network data set size to that of the non-networks. This reduced the model performance for prediction of [Oxidation]_CC_ levels (R^2^ = 0.73), but not to the non-predictable level of the non-networks (R^2^ ≤ 0.1), **(Fig. 2L).** This demonstrates that the combinatorial structural information in the networked mitochondrial components leads to higher predictability of their redox and its intra-network heterogeneity.

Thus, hierarchical mito-SinComp analyses reveal remarkable intra-cellular and intra-network heterogeneity of mitochondrial redox, which can be predicted only for networks by ML based modelling of combinatorial network structural features. Such predictability can be tested for any mitochondrial function and can be potentially utilized in translational settings.

### Use of mito-SinComp structural feature, CRL-mitoNet, reveals conversion of Hyperfused-Mitochondrial-Networks to Small-Mitochondrial-Networks upon stem cell activation

We tested if untangling mitochondrial heterogeneity can reconcile controversies on fragmented mitochondrial entities in stem cells^21,24,25^. We used mito-SinComp to identify the characteristic mitochondrial form of stem cells in benchmarked HaCaT keratinocytes populations where genetic manipulation of Drp1 enriched or depleted Krt15^hi^ stem cells ^23^, and in flow sorted Aldh^hi^ ovarian cancer tumor initiating (stem) cells where Drp1 modulates Aldh expression^21,42^.

To capture both inter- and intracellular mitochondrial heterogeneity in a single feature, we computed the ‘Cell-Relative Length of mitochondrial network or non-network’, [CRL mito-Net] or [CRL mito-nonNet] (**Suppl Table 1**). Comparison of distribution of [CRL mito-Net] in the INT-cells and the HF-cells groups of the Drp1 manipulated keratinocytes, indicate that the INT-cells may be enriched with ‘Small Mitochondrial Networks’ (**SMNs**), whereas the HF-cells with ‘Hyperfused Mitochondrial Networks’ (**HMNs**) and ‘very Small Mitochondrial Networks’ (**vSMNs**)(the smallest network unit with 4 nodes) (**Fig. 3A, Suppl Table 2**). Indeed, classification based on [CRL mito-Net] thresholds (see Methods) confirmed remarkable SMN enrichment in the INT-cells and the enrichment of the HMNs and vSMNs in the HF-cells (**Fig. 3B**); examples of networks and non-networks (which may include the mitochondria derived vesicles ^43^) are shown (**Fig. 3C)**. Mito-SinComp (S) features reveal that in comparison to the HMNs, the SMNs are (**Fig. 3D, Fig. S3A**): **a)** ∼10 fold smaller in size and in abundance of nodes and edges. **b)** comparable in element size within a network; **c)** remarkably heterogenous in thickness, network complexity (nodes or edge density) and maximum connectivity.

**Figure. 3:**
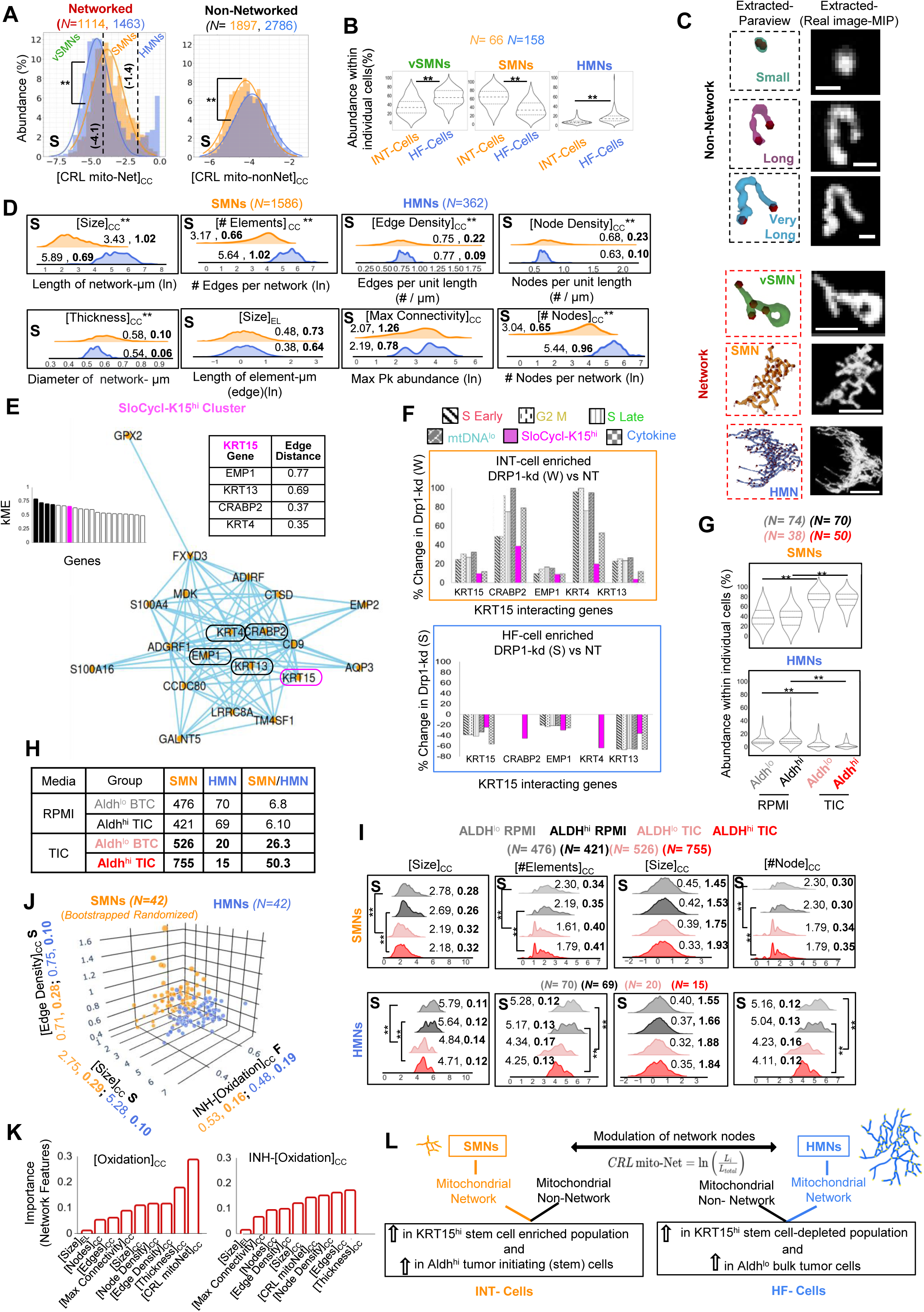
**(A)**. Histogram distribution of [CRL mito-Net] or [CRL mito-nonNet], with demarcation of thresholds (dashed lines) for vSMNs, SMNs and HMNs. (threshold calculation in Methods). **(B)**. Violin plot of vSMNs, SMNs, HMNs abundance in individual cells in color-coded cell groups. **(C)**. Magnified views of the extracted Networks and non-Networks from representative cells (Fig. 2A) with their corresponding micrographs. **(D)**. Ridgeline plots comparing the distribution of the named (S) features between SMNs and HMNs; Median and Coefficient of Variation (in bold) are indicated for each feature. **(E)**. Hub-gene network plot displaying the top 20 genes from the outlined module in **Fig.S3C**; with kME module plot and table (insets) highlighting KRT15 direct interacting genes. **(F)**. Bar plots of percentage change in median expression of KRT15 and its direct interacting partners (from E) in the DRP1 manipulated cells from NT control, across identified cell clusters. **(G)**. Violin plot of SMN and HMN abundance within individual cells of color-coded cell groups. **(H)**. Table showing number of SMNs, HMNs, and SMN / HMN ratio from (G). **(I)**. Ridgeline plots comparing the distribution of the named (S) features of SMNs and HMNs from (G); Median and Coefficient of Variation (in bold) are indicated for each feature. **(J)**. Trivariate S-F scatter plot of named SMN or HMN S-F features identified from Fig. 2D. **(K)**. Bar plot of feature importance values for individual network features from ML based predictive model described in Fig. 2I-L. **(L)**. Schematic describing connection of [CRL mito-Net] classified SMNs and HMNs to KRT15 marked stemness transcriptomic network in Drp1 manipulated cells. **signifies Holmes-Bonferroni corrected p-value < 0.05; Scale bar: 5 μm for Networks and 1μm for non-Networks; N = sample size.

KRT15^hi^ stem cells are enriched by the weak (W) Drp1 knockdown that sustains INT-cells, and depleted due to the strong (S) Drp1 knockdown (kd) that sustains HF-cells ^23^. Therefore, we used this benchmark data set and its corresponding scRNA-seq data to investigate any link of the SMN enrichment to the transcriptomic network of the lineage specific stemness gene KRT15 ^44^. The hdWGCNA **(Fig. S3B-C)** of the combined differentially expressed gene sets from pseudobulk analyses of NT vs Drp1-kd(W) and NT vs Drp1-kd (S) identified the 10 genes that directly interact with KRT15 in its transcriptomic network **(Fig. S3D-F).** HdWGCNA of the marker genes of the KRT15^hi^ stem cell cluster, confirmed 4 out of the above 10 genes as direct KRT15 interacting partners **(Fig. 3E** with kME bar plot and edge distance table as insets, **Fig. S3E-F)**. Importantly, these KRT15 interacting genes undergo marked upregulation across all cell clusters in the KRT15^hi^ stem cell enriched Drp1-kd (W) population with abundant SMNs in the INT subpopulation **(Fig. 3F, top, Fig. S3G)**. On the other hand, two of those genes undergo marked downregulation only in the KRT15^hi^ cells that are depleted in the Drp1-kd (S) population with depletion of SMNs in the HF-cells **(Fig. 3F, bottom)**. Thus, parallel mito-SinComp and hdWGCNA analyses indicate that the SMNs are linked to KRT15-stemness transcriptomic network in KRT15^hi^ stem cells.

To address if activating stemness can enrich SMNs we performed mito-SinComp structural analyses in sorted tumor initiating (stem) cells (TICs) and bulk tumor cells (BTCs) from A2780-CP lines. Self-renewal was activated in the sorted Aldh^hi^ TICs and Aldh^lo^ BTCs with TIC media (24 hrs), using RPMI control media ^21^. The mito-SinComp 3D maps and structural analyses reveal that the TIC media (with growth factors) reduces the average length of networked mitochondria while increasing their abundance in both Aldh^hi^ TICs and Aldh^lo^ BTCs **(Fig. S4A-C)**. TIC media causes a pronounced increase in [CRL mito-Net] in both Aldh^hi^ TICs and Aldh^lo^ BTCs **(Fig. S4D)**, and induces remarkable enrichment of SMNs and a concomitant depletion of the HMNs in them **(Fig. 3G).** More importantly, SMN / HMN ratio indicates that enrichment of SMNs by TIC media is 2 folds higher in the Aldh^hi^ TICs than in the Aldh^lo^ BTCs **(Fig. 3H)**. These SMNs remain ∼10X smaller that of the HMNs and remarkably heterogenous **(Fig. 3I, Fig. S4E**, similar to **Fig. 3D)**, Notably, the TIC media dramatically reduces the network size and abundance of nodes and edges of the HMNs almost to the level of the SMNs, with no impact on individual element size **(Fig. 3I)**. TIC medium increases thickness in both SMNs and HMNs, while reducing maximum nodal connectivity in the SMNs **(Fig. S4E-F)**. Together, these data demonstrate that TIC media converts all the HMNs to SMNs by severing specific nodal connections of the HMNs and not by reducing the element size.

Next, we examined the heterogenous SMNs and the HMNs in the pooled data set where mito-SinComp S-F analyses revealed profound intra-network heterogeneity of [Oxidation]_CC_ (**Fig.2**). Trivariate S-F plot of INH-[Oxidation]_CC_, [Size]_CC_ and [Edge Density] reveal that the SMNs maintain similar intra-network heterogeneity as the HMNs while higher network complexity is exhibited in some SMNs with lower intra-network heterogeneity (**Fig. 3J**). Nonetheless, the [CRL mito-Net], which distinguishes SMNs from the HMNs, accounts for ∼30% of the predictability for redox in the ML modelling, while combination of various structural features predicts intra-network heterogeneity (**Fig. 3K**, **Fig. 2**); expectedly, the invariant [Element Size] accounts for only ∼1% of both predictability.

In summary, we demonstrate that the quantitative feature, [CRL mito-Net], untangles the SMNs from the HMNs and contributes substantially in prediction of network redox status. Mechanistic analyses with mito-SinComp demonstrate that activation of stemness converts the HMNs to ∼10 folds smaller heterogenous SMNs by severing specific HMN nodal connections (**Fig. 3L**). Such precise regulation happens more pronouncedly in the TIC enriched population in the early during self-renewal induction, linking SMN formation to stemness specification.

### Mito-SinComp S-F analyses reveals prompt conversion of oxidized HMNs to redox-tuned SMNs of restricted complexity in non-transformed cells upon exposure to stem cell enriching carcinogen

We used mito-SinComp S-F analyses to directly address if any functionally relevant SMN enrichment is triggered in non-neoplastic HaCaT keratinocytes at the initial stage of stem cell driven transformation. Thus, we studied the acute impact of reported stem cell enriching TCDD-1nM, with control TCDD-10nM carcinogenic dose (with reduced stem cell enrichment) and the Toluene vehicle control ^23^. Interestingly, TCDD-1nM promptly depolarizes mitochondria within one hour (T1-1h) when TCDD-10nM needs 4 hours to cause the same effect **(Fig. 4A, Fig. S5A)**. Mitochondrial depolarization alters their shape ^45^ and redox function ^46^. Therefore, we used mito-SinComp S(network)-F(redox) analyses to test if stem cell enriching T1-1h induced mitochondrial depolarization is linked to formation of SMNs with specific redox functions using the Toluene, T1-1h and T10-1h groups from the experimental pool of **Fig. 2**.

**Figure. 4:**
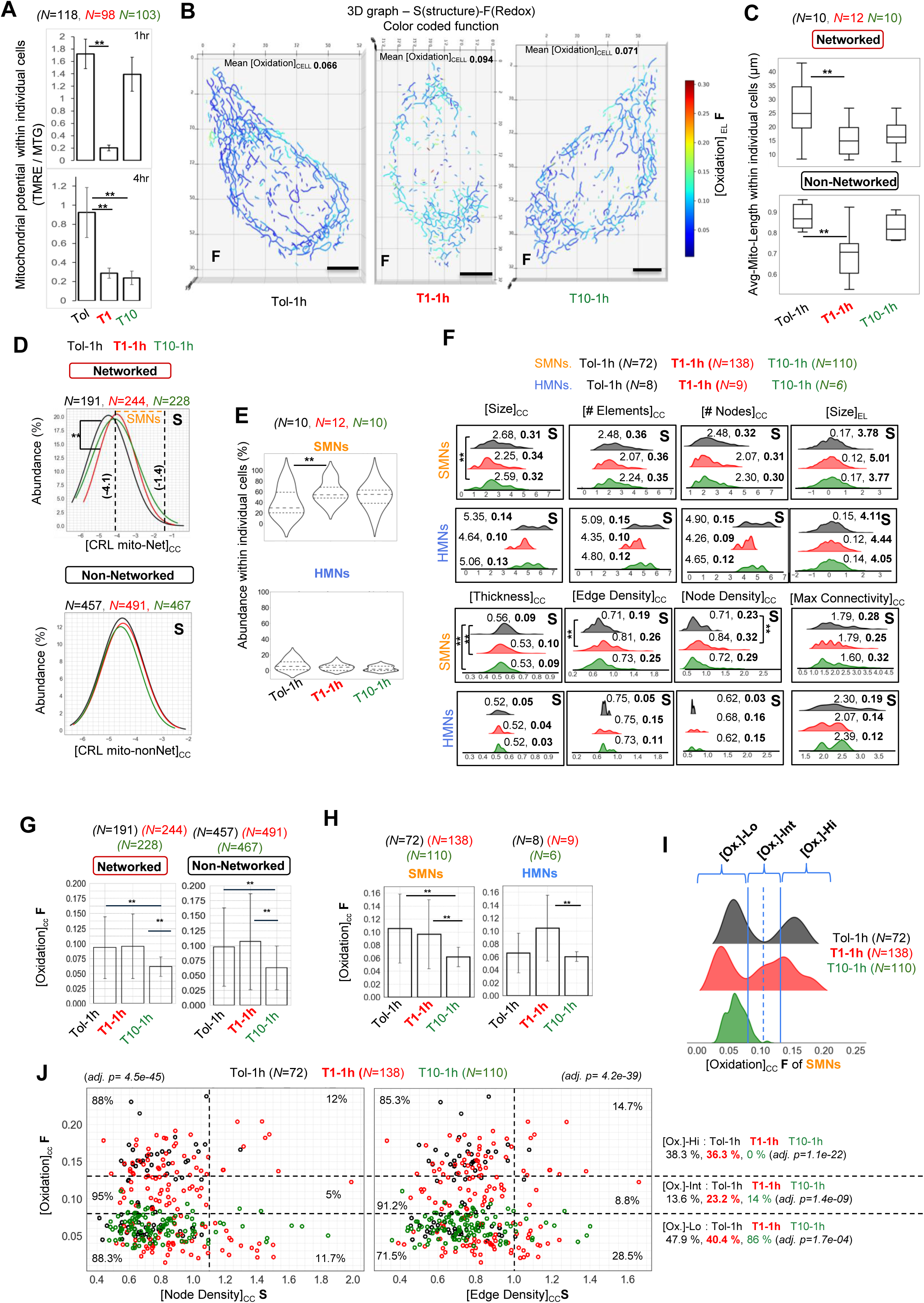
**(A)**. Mitochondrial potential of individual cells measured from confocal micrographs of individual HaCaT cells exposed to Toluene, TCDD 1nM or 10nM for 1 or 4 hours. **(B)**. Representative 3D S-F Graphs of HaCaT cells stably expressing mito-PSMO2 and mito-roGFP from the labelled groups showing [Oxidation]_EL_ with colour-coded scale. **(C)**. Box plot of average length mitochondrial Networks and non-Networks within individual cells in labelled groups. **(D)**. Histogram distribution of [CRL mito-Net] or [CRL mito-nonNet] in labelled groups with demarcation of thresholds (dashed lines) for SMNs and HMNs (as in Fig. 3A). **(E)**. Violin plot of SMN and HMN abundance within individual cells in labelled cell groups. **(F)**. Ridgeline plots of distribution of the named (S) features of SMNs and HMNs between labelled cell groups; Median and Coefficient of Variation (in bold) are indicated for each feature. **(G)**. Bar plots of mean of [Oxidation]_CC_ of Networked and non-Networked mitochondria of labelled groups; errors: SD. **(H)**. Bar plots of [Oxidation]_CC_ in SMNs and HMNs in the color coded groups; errors: SD. **(I)**. Ridgeline plot of the distribution of [Oxidation]_CC_ in SMNs in the color coded groups. The solid and dashed lines distinguish the [Ox.]-Hi, [Ox.]-Int and Ox-Lo, where the solid line demarcates minimum abundance between the two peaks of Tol-1h control and dashed lines on both sides demarcates the distance to the upper quartile level of the T10-1h control. **(J)**. Bivariate S-F scatter plots of [Oxidation]_CC_ and [Edge or Node Density] for SMNs in the color coded groups; vertical and horizontal lines demarcate the S and F features, respectively. **signifies Holmes-Bonferroni corrected p-value < 0.05; Scale bar: 5 μm; N = sample size

The 3D S(structure)-F(redox) maps of representative cells indicate lack of large mitochondrial networks particularly after T1-1h exposure, similar to the TIC media, and elevated [Oxidation]_CELL_ **(Fig. 4B, S5A-B)**. This is quantitatively reflected in reduced average length of both networked and non-networked mitochondria likely due to the overall increased contribution of mitochondrial fission, which is modest with T10-1h exposure (**Fig. 4C, S5C-F)**. [CRL mito-Net] based classification and analyses of various structural features confirms that the stem cell enriching T1-1h dose promptly converts all the HMNs to 10X smaller and heterogenous SMNs by severing HMN nodal connections, while T10-1h exposure exhibits less specificity of SMN enrichment **(Fig. 4D-F, Fig. S5G)**. Although this is similar to stem cell activation with TIC medium **(Fig. 3G-I)**, a notable difference is that T1-1h exposure elevates network complexity (node or edge density) **(Fig. 4F)**.

The pronounced specific impact of T1-1h on mitochondrial potential **(Fig. 4A)** and on the HMN structure **(Fig. 4F)** associates with significant increase in [Oxidation]_CC_ only in the HMNs that are converting to SMNs, which remain undetected in the uncategorized networked mitochondria **(Fig. 4G-H)**. On the other hand, the [Oxidation]_CC_ is bimodally distributed in the SMNs of the Tol-1h and T1-1h groups and reduced to a unimodal distribution in the T10-1h group (**Fig. 4I**). To tease out such heterogeneity of SMN oxidation, we studied the structural parameters in [Ox]_CC_ -Hi / Int / Lo zones **(Fig. 4I**, zone demarcations are defined in the legend). T1-1h exposure reduces SMN size and elevates their network complexity prominently in the [Ox]_CC_ -Lo zones (**Fig. S5H**). To probe further, we performed bivariate S-F analyses between [Oxidation] and individual structural parameters of the SMNs and quantified their abundance in the defined [Oxidation]_CC_ zones. The [Ox]_CC_ -Int zone is overall enriched (∼1.7 X) with T1-1h induced SMNs and [Ox]_CC_-Lo zone is enriched (∼2X) with T10-1h induced SMNs, with respect to the Tol-1h group **(Fig. 4J)**. Interestingly, the bivariate S-F plot between [Node or Edge density] and [Oxidation]_CC_ appears to be unique from other structural parameters **(Fig. S5I)**. Particularly, the SMNs are primarily restricted within a lower level of [Node or Edge density] and maximal nodal connectivity only in the [Ox]_CC_ -Int zone **(Fig. 4J, Fig. S5I)**.

Thus, mito-SinComp S-F analyses demonstrate that exposure of non-transformed cells to the stem cell enriching carcinogen promptly oxidizes their HMNs and severs nodal connections to convert them to the heterogeneous SMNs that are prevented from conversion to the vSMNs. Importantly, a subpopulation of SMNs remain tuned at an intermediate redox level and with restricted network complexity. Such unprecedented untangling of mitochondrial structure-function heterogeneity can be achieved by mito-SinComp in any cell physiology.

### Carcinogen induced enrichment of redox-tuned SMNs specifies stemness by establishing mtDNA-stemness transcriptomic network, revealed by coupled scRNA-seq

We employed scRNA-seq to investigate the immediate cellular impact of T1-1h induced enrichment of redox-tuned SMNs elucidated by mito-SinComp. Therefore, we performed scRNA-seq of HaCaT cells following SMN enrichment i.e. after 2.5 hrs exposure to Tol (vehicle control), T1 (stem cell enriching dose) and T10 (control dose).

Pairwise pseudo-bulk scRNA-seq analyses confirmed upregulation of the canonical TCDD target gene, *CYP1B1* ^47^ in T10-2.5h but not in T1-2.5h (not shown). Surprisingly, various protein coding genes encoded by the mtDNA heavy strand are upregulated presumably in the SMNs that are the predominant mitochondrial form maintained by T1-2.5h exposure **(Fig. 5A)**; the *MT-ND6* gene product coded by the mtDNA light strand is downregulated only by T10-2.5h.

**Figure. 5:**
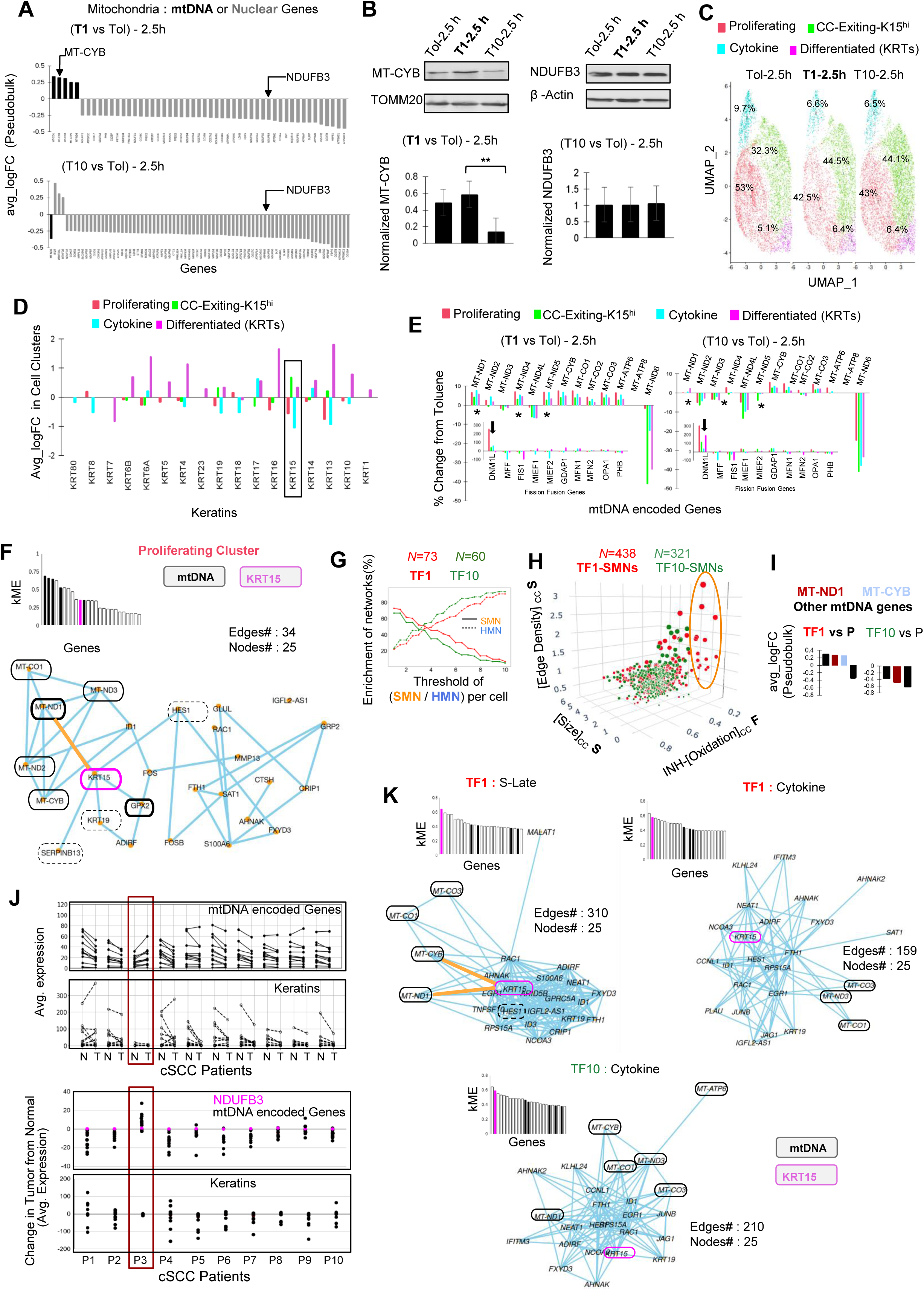
**(A)**. Pairwise pseudo-bulk analyses of genes encoding mitochondrial proteins (mtDNA or Nuclear DNA encoded) from scRNA-seq data of HaCAT cells in labelled groups. **(B).** Immunoblot analysis of MT-CYB and NDUFB3, with TOM20 and β-actin as loading controls in labelled groups; quantified levels are normalized by loading control. **(C).** UMAP plot of scRNA-seq derived clusters within cell populations of the labelled groups; the numbers indicate % abundance of cells in each cluster based on Fig. S5C (see Methods for functional annotation). **(D).** Bar plots showing average logFC of all Keratin genes in the named cell clusters, with stem cell-specific KRT15 boxed; (Krt15 raw data in Fig. **S8A**). **(E).** Bar plots showing the percentage change in median expression of mtDNA-encoded genes relative to the control in labeled groups across cell clusters (raw data in **Fig. S8B**); * mark MT-ND1, MT-ND4, MT-ND5; inset shows bar plots with for the core fission–fusion genes, with arrows on DNM1L gene for Drp1. **(F).** Hub-gene network plot displaying the top 25 genes from the kME plot (inset) highlighting KRT15 interacting mtDNA genes. **(G).** Line plot showing percentage enrichment of SMNs and HMNs along the threshold values of SMN / HMN ratio within individual cells in the labelled groups. **(H).** Trivariate S-F scatter plot of named (S) and (F) features for networked mitochondrial components in color coded groups. **(I).** Bar plot showing differential expression of mtDNA genes in the labelled groups. **(J).** Absolute average expression of mtDNA genes and major keratins in cSCC patients (top) in normal (N) and tumor (T) tissue samples (top), while change in average expression in tumor from normal (bottom); original data from reference 50. **(K).** Hub-Gene Network plot displaying the top 25 genes from the kME plot (inset) highlighting KRT15 interacting mtDNA genes.

This distinction between the mt-DNA heavy and light stand suggests a distinct transcriptional regulation from the distinct promoters at each stand^48^. In contrast, the nuclear genes encoding mitochondrial proteins are overall downregulated by both T1-2.5h and T10-2.5h **(Fig. 5A)**. Immunoblot analyses validated the contrasting effects of T1-2.5h and T10-2.5h on mt-DNA coded MT-CYB levels, with no impact on nuclear DNA coded NDUFB3 **(Fig. 5B)**. GSEA based functional annotation of genome wide pairwise pseudo-bulk analyses demonstrate that both T1-2.5h or T10-2.5h downregulate genes involved in mitochondrial energetics and cell cycle **(Fig. S6A)**. Furthermore, hdWGCNA demonstrates that the modules obtained with (T1 vs Tol)-2.5h gene sets have higher number of mito-Energetics and CellCycle genes, and their transcriptomic connectivity is strikingly complex compared to the similar module identified with (T10 vs Tol)-2.5h gene sets (**Fig. S6B-D)**.

Next, we identified the cell clusters on a UMAP plot, quantified their abundance **(Fig.5C, S6E)**, and assigned cluster functionality by GSEA on their respective gene markers **(Fig. S6F)**. Both T1-2.5h or T10-2.5h reduces abundance of ‘Proliferating’ cells (upregulated cell cycle makers) and increases abundance of the ‘CC-Exiting’ cells (downregulated cell cycle makers) (**Fig.5C)**. We re-designated the latter as ‘CC-Exiting-K15^hi^’ due to highest contribution of the stem cell gene KRT15 (among 18 KRT genes) in defining this cluster **(Fig. 5D, Fig.S7A)**.

Noteworthily, the specific T1-2.5h driven upregulation of mtDNA heavy strand genes happens across all cell clusters and most prominently for the components of the major site of mitochondrial ROS production namely Complex I ^46^ (marked by * in **Fig. 5E**, outlined in **Fig. S7B)**. This suggests a direct impact of T1-2.5h on mtDNA gene expression in the SMNs, although their lower expression in the ‘CC-Exiting-K15^hi^’ cluster is consistent with their reciprocal relationship with KRT15 in the transformed population ^23^. Interestingly, expression of Drp1(coded by DNM1L gene), among key genes modulating mitochondrial fission and fusion, is differentially modulated between T1-2.5h and T10-2.5h (arrows **in Fig. 5E** inset), consistent with Drp1 knockdown modulating stem cell status in this lineage ^23^. Next, we used hdWGCNA to identify any mtDNA-KRT15 (stemness) interacting modules within each cell cluster and further assigned functionality to the listed modules using GSEA. Importantly, the mtDNA genes are found to interact with the stem-cell gene KRT15 only in the Proliferating cluster of the T1-2.5h group (kME bar plot as inset in **Fig. 5F, Fig. S7C-D**). Hub-gene network analyses of the mtDNA-KRT15(stemness) module reveal a distinct network of interacting mtDNA genes, out of which redox related MT-ND1 of Complex I is directly connected to KRT15 that is further connected to the major redox regulator GPX2 **(Fig. 5F)**.

To confirm the above findings in neoplastic stem cells, we performed parallel mito-SinComp and hdWGCNA on TCDD-1nM transformed experimental group with pronounced stem cell enrichment (TF1), TCDD-10 nM transformed control group with substantially weaker stem cell enrichment (TF10) and non-transformed parental control group (P) ^23^. [CRL mito-Net] based network classification demonstrates that SMN and HMN abundance is distinctly different between the TF1 and the control groups **(Fig. S7E-F)**. The percentage of cells above or below a given threshold of SMN / HMN ratio in each cell signifies SMN or HMN enrichment, respectively. Indeed, SMN enrichment remains consistently higher in the TF1 population between a threshold of 4 and 6 (**Fig. 5G, Fig. S7G)**, while SMNs and HMNs maintain their distinctive properties (**Fig. S7H**). Moreover, trivariate mito-SinComp S-F analyses demonstrated that the SMNs that exhibit maximum heterogeneity of [Edge Density]_CC_ and remarkably low INH-[Oxidation]_CC_ belong to the stem cell enriched TF1 population (outlined in **Fig. 5H**).

Probing into the mt-DNA gene expression confirmed specific upregulation of redox related MT-ND1 gene for Complex I and MT-CYB gene for Complex III particularly in TF1 **(Fig. 5I, Fig. S7I)**. Also, specifically TF1 group exhibits prominent downregulation of the mt-DNA regulating molecule Prohibitin particularly in the previously labelled KRT15^hi^ stem cell cluster (**Fig. S7I**, empty triangle, **Fig. S7J**)^49^. Next, we confirmed the SMN related distinction of mtDNA genes in the available scRNA-seq data from the relevant cutaneous squamous cell carcinoma (cSCC) patient samples ^50^. Interestingly, 1 out of 9 patients exhibits prominent increase in mtDNA genes (including MT-ND1 and MT-CYB) in the cSCC tumor from the adjacent normal tissues, similar to the SMN and stem cell enriched TF1 cells **(Fig. 5J, outlined)**. Also, this patient exhibits no alteration in keratin gene expression in contrast to the other patients showing decrease in mtDNA genes (similar to TF10) and keratin genes in the tumor. Expectedly, the nuclear encoded gene for the mitochondrial protein NDUFB3 shows no change across all the 10 patients. Finally, hdWGCNA confirmed the direct interaction of KRT15 with MT-ND1 (and MT-CB5) in a module detected particularly in the S-Late cluster of the TF1 **(Fig. 5K** with kME plots as inset**, Fig. S7K)**; although the Cytokine clusters of the TF1 and TF10 populations harbor a similar module, no direct MT-ND1-KRT15 connection was detected.

In summary, coupled hdWCGNA demonstrated that enrichment of the redox-tuned SMNs by the stem cell enriching carcinogen is promptly followed by upregulation of their mtDNA heavy strand genes and establishment of the MT-ND1(redox)-KRT15(stemness) transcriptomic network in the proliferating non-transformed cells. Given the same features are detected only in stem cell enriched transformed population, we conclude that SMN induced transcriptomic modulation prime stemness driven neoplasticity in non-transformed cells. Identification of cSCC patient with SMN linked mtDNA changes indicate the clinical relevance of our finding while highlighting patient heterogeneity.

### Mito-SinComp analyses of mtDNA nucleoid organization in fixed samples demonstrates higher nucleoid abundance in the characteristic stemness specifying SMNs

Efforts are underway to quantitatively resolve the remarkable heterogeneity of the polyploid mtDNA that is organized in nucleoids of variable size and number within mitochondrial networks ^51^. Using mito-SinComp modular design **(Fig. 1),** we investigated if the mechanism of increase in mtDNA gene expression with conversion of HMNs to SMNs involves functional organization of their mtDNA. We performed in-depth mito-SinComp S-F analyses in HaCaT keratinocyte and beyond cell boundaries in a *Drosophila* tissue amenable to confocal microscopy, using nucleoid number and size as functional surrogates.

The individual mtDNA nucleoids are identified from the high-resolution 3D confocal micrographs as pixel clusters using DBSCAN algorithm after appropriate optimization of its hyperparameters (see Methods) (**Fig. 6A**). The identified clusters are mapped to the corresponding mitochondrial structural coordinates and mitochondrial components, where each component can harbor multiple mtDNA nucleoids of varying size (arrows in **Fig. 6A)**. The visually confirmed unmapped clusters (arrowheads in **Fig. 6A)** may represent released mtDNA nucleoids ^52^ or other unwanted signal. Fixed mito-SinComp analyses confirm the higher heterogeneity in [Thickness]_CC_ of the non-networked components **(Fig. 6B)**, as in live cells (**Fig. 2E)**. However, the absolute [Size]_CC_ and [Overall Nodal Connectivity] are reduced in fixed cells (**Fig. S8A,B**). We computed the following mtDNA nucleoid functional features for individual mitochondrial components: [Nucleoid Number]_CC_, [Nucleoid Density]_CC_, [Nucleoid Size]_EL_, [Avg-Nucleoid Size]_CC_ and INH-[Nucleoid Size]_CC_ (**Suppl. Table 1**). The networked components have higher [mtDNA nucleoid Number]_CC_ but reduced [Nucleoid Density]_CC_ in comparison to the non-networked components, while [Nucleoid Size]_EL_ remains comparable **(Fig 6B)**.

**Figure. 6:**
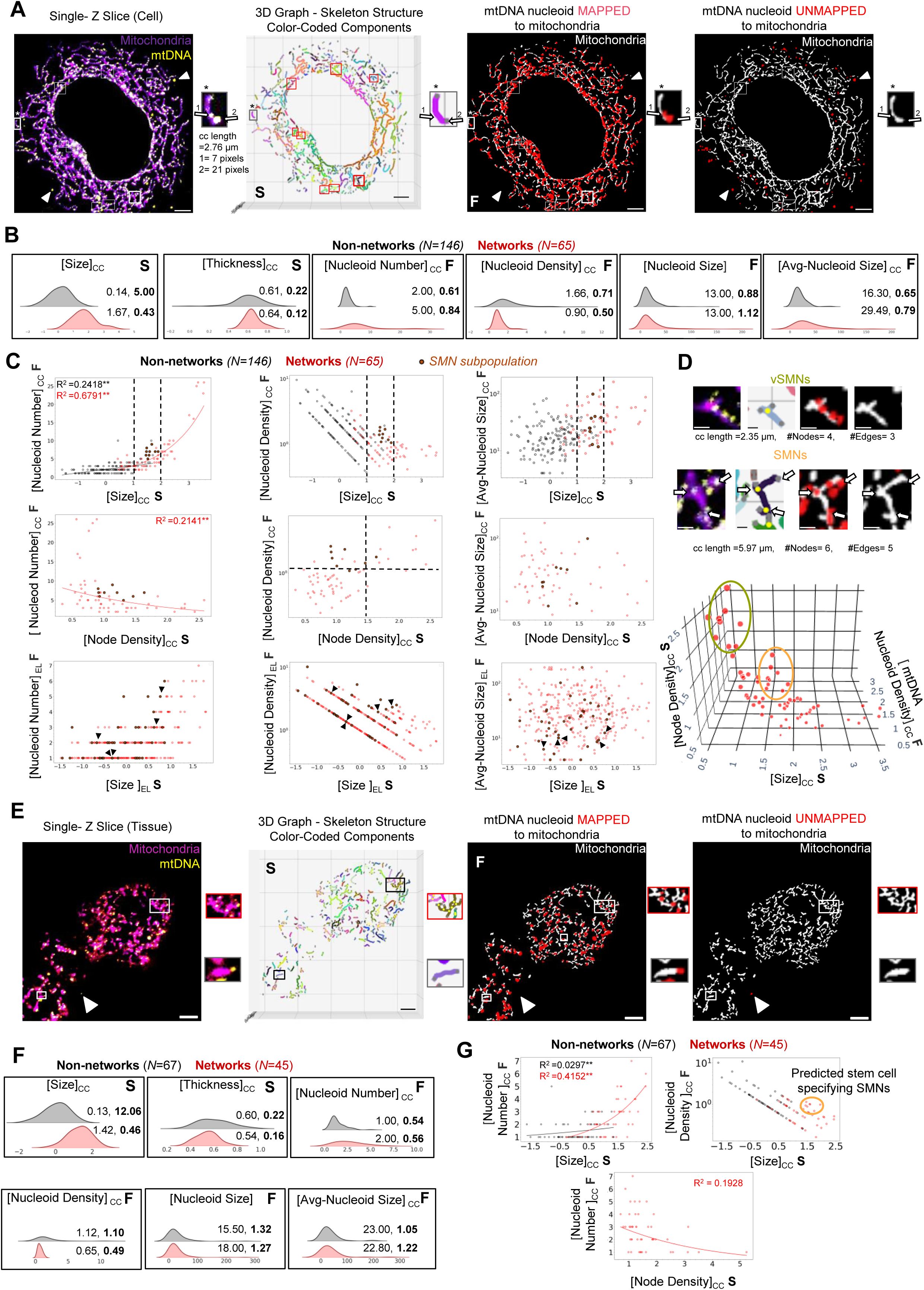
Quantitative mito-SinComp S-F analyses of functional organization of mtDNA within networks and non-networks in fixed cells and tissues. **(A)** Representative micrograph of a co-immunostained HaCaT cell with its 3D Structure Graph and Mapped / unmapped mtDNA nucleoids projected on MitoGraph binary image; outlined are identified SMNs and magnified non-network (*) with its feature dimensions mentioned. **(B)** Ridgeline plots comparing distribution of the S-F features of mitochondrial Networks and non-Networks; Median and Coefficient of Variation (in bold) are indicated for each feature. **(C)** Bivariate S-F scatter plots for and Networks and non-Networks identified SMNs highlighted. **(D)** Trivariate S-F 3D scatter plot of S-F features for Networks in color coded groups. **(F).** Representative micrograph of co-immunostained *Drosophila* fat body tissue with 3D Structure and the Mapped / unmapped mtDNA nucleoids projected on MitoGraph binary image; representative Networks and non-Network outlined and magnified. **(G).** Ridgeline plots comparing distribution of S-F features of mitochondrial Networks and non-Networks; Median and Coefficient of Variation (in bold) are indicated for each feature. **(H).** Relevant bivariate S-F scatter plots for mitochondrial non-networked and networks; predicted SMN subpopulation outlined. ** signifies p-value < 0.0005; Scale bar: 5 μm; N = sample size.

Interestingly, S-F plot of [Size]_CC_ vs [Nucleoid Number]_CC_ reveal that while longer non-networked mitochondria have upto 5 mtDNA nucleoid in this cell, the [Nucleoid Number]_CC_ exponentially increases after formation of mitochondrial network of a critical window of [Size]_CC_ **(Fig. 6C**, dashed lines). For any given [Nucleoid Number]_CC_, the [Nucleoid Density]_CC_ is expected to reduce with [Size]_CC_. Deviation from this expectation indicates influence of other factors, as is evident in certain networked components in the critical window (highlighted in **Fig. 6C**). These networks are identified as an SMN subpopulation by quantitatively comparing similar S-F plots of [Node Density]_CC_ vs [Size]_CC_ in the fixed cell with that of live cells where SMNs were defined (**Fig. S8B**). Other S-F plots confirm that the higher nucleoid density in these SMN subpopulation is due to their higher nucleoid number. Importantly, their restricted network complexity indicate they may represent the redox-tuned SMNs subpopulation with restricted network complexity (**Fig. 4J**). Like other networks, these SMNs maintain intra-network heterogeneity of nucleoid number and size between their elements (arrowheads in **Fig. 6C** for the example SMN in **Fig. 6D**). Visual examinations reveal the positioning of nucleoids at the nodes of these SMNs (arrows in **Fig. 6D, top**). A trivariate plot of [Nucleoid Density]_CC_, [Node Density]_CC_ and [Size]_CC_ distinguishes these SMN from the VSMs which exhibit maximized [Nucleoid Density] and [Node Density] due to their small size (**Fig. 6D, bottom**). Mito-SinComp analyses in *Drosophila* fixed tissue beyond cell boundaries recapitulates the key distinction of mtDNA organization between networks and non-networks (**Fig. 6E-G**), highlighting fundamental importance of our findings. Notably, the S-F plot of [Size]_CC_ and [Nucleoid Density] identifies the potential SMN subpopulation which when enriched by neoplastic insults may specify stemness in the cells carrying these SMNs, thus priming neoplasticity (outlined in **Fig.6G**).

In summary, mito-SinComp analyses of mtDNA-nucleoids revealed their distinctive organization in the networks and non-networks, and also their positioning at the internal and free nodes. Comparative mito-SinComp analyses between fixed cells and in live cells, in conjunction with scRNA-seq, reveals a novel mechanistic model of stem cell specification (**Fig. 7**). Stemness specification involves enrichment of a subpopulation of redox-tuned SMNs with restricted network complexity and greater mtDNA abundance, generated by precise breakdown of oxidized HMNs. This supports the elevated mtDNA gene expression in these SMNs, which in turn establishes the transcriptomic link between mtDNA coded mt-ND1 (redox relevant) gene and nuclear DNA coded Krt15 (stemness gene).

**Figure 7:**
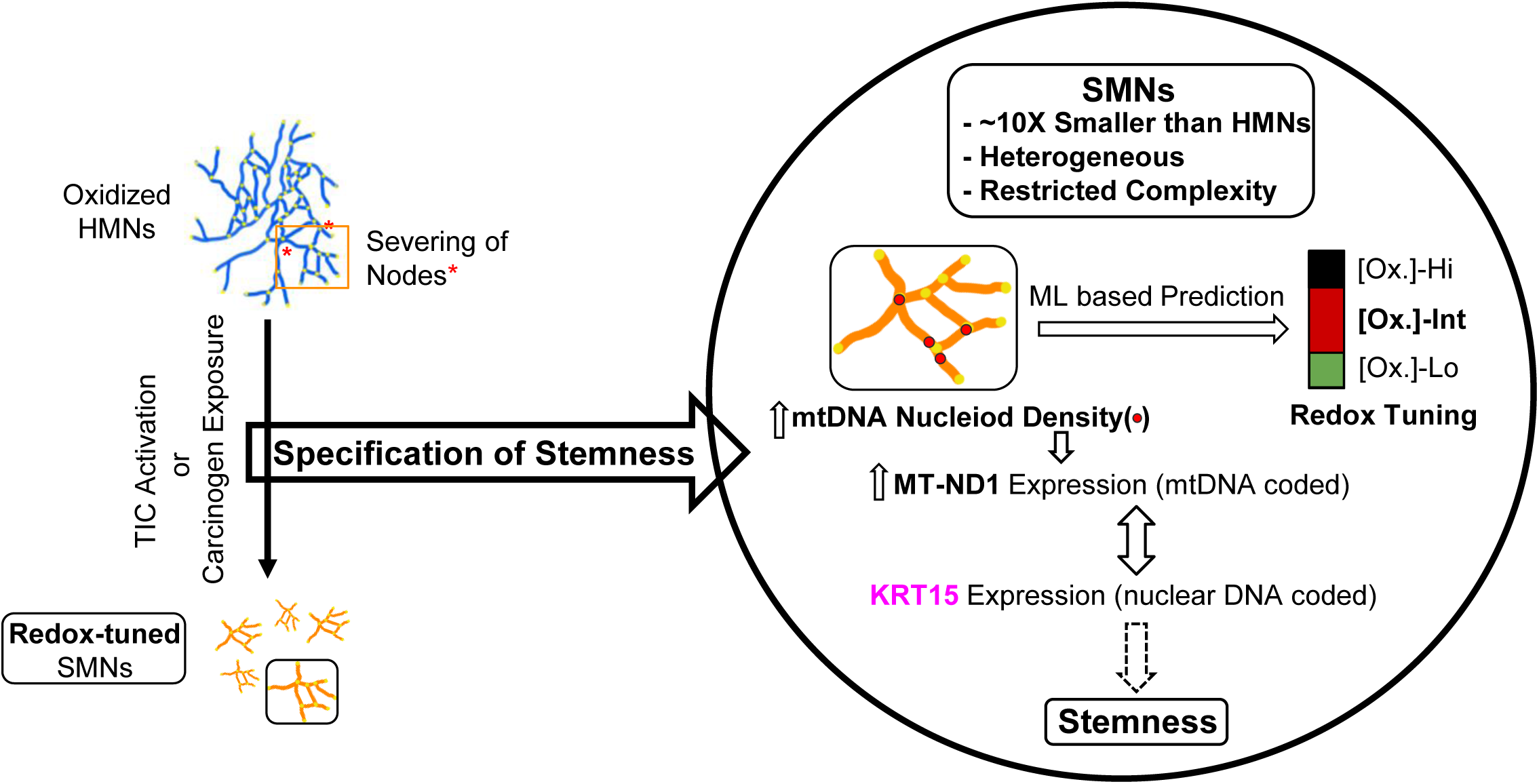
Schematic illustration of stem cell specification by SMNs. The illustration summarizes the mechanism of stemness specification as revealed by mito-SinComp S-F analyses and molecular players identification by coupled scRNA-seq analyses. Stemness specification induces severing of specific nodes of oxidized HMN (*) to convert them to characteristic redox-tuned SMNs (outlined) where elevated mt-DNA nucleoid density supports increased mtDNA gene expression. This establishes the MT-ND1(redox)-KRT15(stemness) transcriptomic network.

## Discussion

Quantitative understanding of mitochondrial heterogeneity is required to elucidate the complex role of mitochondria in normal and pathophysiological conditions^1,5,53–55^. Understanding heterogeneity requires quantitative analyses of single entities, which is achieved by Mito-SinComp structure(S)-function(F) analyses (**Fig. 1**). Mito-SinComp S and F analyses focused on redox function revealed redox levels and its remarkable intra-network redox heterogeneity can be quantitively predicted by a combination of the structural features through a machine-learning approach (**Fig. 2,3**). Mito-SinComp analyses suggest that a characteristic mitochondrial form, SMNs with specific redox and mtDNA properties, specifies stemness. Our conclusions are based on mito-SinComp data from 17 experimental groups, involving 772 cells, 12181 networks, 24556 non-networks, 848 HMNs and 5872 SMNs, across 7 cell lines (parental and derived by genetic manipulation or carcinogen exposure). Moreover, using publicly available data ^50^, we demonstrate inter-cancer patient heterogeneity for SMN linked mtDNA modulation, (**Fig. 5J**). Furthermore, using *Drosophila* we demonstrate that the characteristic SMNs can be identified using mito-SinComp in normal fixed tissues that are amenable for confocal microscopy (**Fig. 6E-G**). Molecular surrogates for SMNs remain to be discovered, which can be used in tissues that are not-amenable for microscopy. Nonetheless, our findings highlight that stem cell functionality of mitochondria may be exhibited not by fully fragmented mitochondria but rather by characteristic SMNs which are presumably generated by precisely regulated fission events and may be maintained by precisely regulated fusion events. This explains need of both fission and fusion processes in stemness, thus reconciling various contradictory results ^3,21,23–25,31–34^.

The unprecedented abilities of mito-SinComp analyses identified and characterized the subpopulation of stemness specifying SMNs, as well as revealed mechanism of their formation and function in stemness specification **(Fig. 7)**. The stemness specifying SMNs are 10 times smaller than HMNs, are redox tuned, have restricted network complexity and elevated mtDNA nucleoid density. This supports elevated mtDNA gene expression and the resulting transcript of the redox relevant mt-ND1 gene maintains direct transcriptomic connection with lineage specific stemness gene, as revealed by coupled hdWGCNA of scRNA-seq data. The redox-tuned SMNs are formed by severing of the nodes of the oxidized HMNs, likely mitochondrial fission (**Fig. S5F**). We speculate that inhibition of HMNs to SMNs conversion by obliteration of mitochondrial fission underlies its marked inhibition of oncogene induced neoplastic transformation ^23,35,36^, which can also be tested in other models of neoplastic transformation.

The SMNs need to be sustained further by keeping mitochondrial fission in check, which explains the fine-tuned regulation of Drp1 in promoting neoplasticity ^23^. Mitochondria act as a hub of redox signaling, while stem cells are particularly dependent on redox tuning ^2,40^. SMN formation could be a potential spatio-temporally controlled mechanism of fine-tuning redox signaling through mitohormesis ^56^. This conception adds a novel direction in elucidation of the bidirectional crosstalk between mitochondrial structure and redox properties ^41,57,58^. hdWGNCA detected mt-ND1(redox)-KRT15(stemness) transcriptomic network in stemness primed proliferating cells and in late S phase after transformation (**Fig. 5**). Therefore, the stemness specification by redox driven SMNs might involve cell cycle regulation, which plays a deterministic role in stemness ^40,59^.

We demonstrated the following capabilities of the quantitative mito-SinComp approach, that can be used for any function of mitochondria for any cell physiology in live and fixed cells and tissues: **A)** distinguishing mitochondrial networks from non-networks, and their properties; **B)** distinguishing component types using quantitative [CRL mito-Net / non-net] feature, and subpopulation of types based on their function; **C)** understanding query driven quantitative S-S, S-F or F-F relationships (bivariate or multivariate) within identified types and subtypes; **D)** quantitative elucidation of intra-network heterogeneity of function; **E)** comparison of quantitative features between live and fixed samples; **F)** usefulness of S-F features in machine-learning based predictive modeling for translational research; **G)** distinguishing impact of agents on specific kinds of networks or non-networks; **H)** informing about mechanism of interconversion between network forms**; I)** elucidating impact of identified components on cell physiology using correlatively or causatively with other quantitative modalities. To further elucidate the molecular mechanism of our novel findings, mito-SinComp can be used in a time-lapse mode to study activation of Drp1 driven fission sites at the HMN nodes, or to identify players involved in maintaining the marked intra-network redox heterogeneity, or to link functional mtDNA organization to their replication, transcription, inheritance etc. Using mito-SinComp on primary cells from patients will enable studying inter-patient and intra-tumor heterogeneity of SMNs towards personalized treatment ^29,60^. Mito-SinComp can be used to study the impact of genetic mutations, recruitment and release of mitochondrial regulators and moonlighting proteins.

In conclusion, we untangled multilevel mitochondrial heterogeneity by directly studying structure-function of single mitochondrial components in the context of stemness using a first of its kind quantitative approach, mito-SinComp. The unprecedented capabilities of mito-SinComp can be used in a variety of ways to make fundamental discoveries and support their translational applications.

## Materials and Methods

### Cell lines

We have used the following parental and derived cell lines across 2 cell lineages: **a)** the parental HaCaT line (immortalized normal skin keratinocytes) and their derived lines transformed with 1nM or 10nM TCDD (**Fig. 5**); **b)** parental HaCaT lines stably expressing weak or strong Drp1 shRNA or non-targeted (NT) shRNA (**Fig. 2,3**); **c)** parental double stable HaCAT line expressing mito-PSmO2 and mito-roGFP and their derived lines transformed with 1nM or 10nM TCDD or 10nM Toluene (**Fig. 2, 4); d)** A2780-CP line (carboplatin resistant ovarian cancer line derived from parental A2780 line) (**Fig. 3**). All HaCaT cells were maintained in Dulbecco’s Modified Eagle’s Medium (DMEM)-high glucose supplemented with 10% FBS and Penicillin-Streptomycin cocktail (100 µg/mL), in a 5% CO_2_ at 37deg C using standard techniques. A2780-CP cells were similarly maintained in RPMI medium supplemented with 1 mM sodium pyruvate.

The double stable HaCaT lines expressing mito-PSmO2 and mito-roGFP were generated using the pCDH lentiviral expression system with puromycin and hygromycin selection, respectively, using standard methods. Transformed HaCaT lines were derived from the Parental HaCaT cells by selecting multiple clones following treatment with 1nM or 10nM dose of TCDD for 16 to 20 days, with media replenishment every 2–3 days. The HaCaT cells with stable knockdown of Drp1 (weak or strong) were generated using Lipofectamine mediated transfection of human DNM1L shRNA1 and shRNA2 with non-targeted control, and were selected using puromycin. All stable genetically modified or transformed cell lines, along with their respective parental controls, were subjected to two consecutive rounds of BM-Cyclin treatment to ensure the removal of any potential mycoplasma contamination. For experiments involving acute exposure of TCDD, the parental or double stable HaCaT cells were exposed to the 1 nM or 10nM TCDD for 1 or 4 hours.

### Cell Sorting using flow cytometry

A2780-CP cells were sorted using flow cytometry based on their ALDH (aldehyde dehydrogenase) activity detected by ALDEFLUOR staining carried out as per the manufacturer’s instructions (Stem Cell Technologies), with minor modifications as required. Stained cells were analyzed using a 488 nm excitation laser with a 525/50 band-pass filter on the LSR II flow cytometer, and a 530/30 band-pass filter on the FACS Aria II, both in combination with a 505 nm long-pass filter. The Aldh activity based cell sorting was performed using BD FACS AriaII. The sorted cells were collected in complete RPMI medium or TIC medium composed of RPMI supplemented with 1× N1 supplement, 500 mg/ml insulin, 20 ng/ml human epidermal growth factor (EGF), and 10 ng/ml basic fibroblast growth factor (bFGF). Following sorting, cells were plated in Geltrex coated LabTek chambers and stained with Mitotracker 633 after 24 hours.

### MitoTracker and TMRE Staining

Cells were incubated in appropriate medium containing MitoTracker Green FM (50–100 nM) or MitoTracker Red FM 633 (250 nM) for 15 minutes at 37 °C. For assessment of mitochondrial membrane potential, TMRE was used at a final concentration of 50 nM under the same incubation conditions. To assess mitochondrial membrane potential relative to mitochondrial mass, cells were first stained with TMRE and then mitotracker green, while TCDD/Toluene doses were maintained in the in the staining and washing media. Stained cells were promptly imaged live on confocal microscope using appropriate acquisition settings, as described below.

### Immunoblotting and Immunofluorescence

For immunoblotting, whole-cell lysates were prepared using Laemmli buffer with reducing agent. Proteins (cell equivalent) were resolved on 10% SDS–polyacrylamide gels and transferred onto PVDF membranes. After blocking with 5% bovine serum albumin (BSA), membranes were incubated with specific primary and then compatible HRP-conjugated secondary antibodies, followed by detection using chemiluminescence, using standard protocols. Densitometric analysis of protein bands was performed using ImageJ, with normalization by loading and experimental controls as appropriate.

For immunofluorescence, cells were cultured on Nunc Lab-Tek chamber slides and processed using standard protocols. Cells were fixed in freshly prepared 4% paraformaldehyde containing 4% (w/v) sucrose, then permeabilized using 0.1% Triton X-100. After permeabilization, cells were blocked in 1% BSA, then stained with specific primary and secondary antibodies. Finally, samples were mounted using Fluoromount-G containing Hoechst 33342 (10 µg/mL) for nuclear counterstaining. Slides were imaged using a confocal microscope as described below. Drosophila fat body tissues were dissected in a glass well plate and immediately fixed in freshly prepared 4% paraformaldehyde. Following fixation, tissues were permeabilized using 0.1% PBS-Triton X-100 (PBS-TX) and subsequently blocked with an appropriate concentration of 10% BSA dissolved in 0.1% PBS-TX. The samples were then incubated with specific primary and compatible secondary antibodies for immunostaining. After antibody incubation, tissues were washed three times with PBS-TX. During the final wash, Hoechst 33342 (1:1,000 dilution) was added to stain nuclear DNA. Finally, the stained fat body tissues were carefully transferred onto a drop of mounting medium on a glass slide. A coverslip was gently placed over the tissues and lightly pressed to achieve an even and uniform spread.

### Confocal Microscopy

Microscopy was performed using either Zeiss LSM 700 laser scanning confocal microscope or Zeiss LSM 900 laser scanning microscope. LSM 700 was equipped with a 40X Plan-Apochromat 1.4 NA oil immersion objective, housed within a temperature- and CO₂-controlled incubation chamber to maintain physiological conditions (for live cell imaging). Appropriate excitation laser lines were used for each fluorophore (405 nm and 488 nm for mito-roGFP, 555 nm for basal mito-PSmO2, 488 nm for MitoTracker Green FM, 633 nm for MitoTracker Red FM 633 and 555 nm for TMRE). Emission was collected using appropriate detectors optimized for each fluorophore. Multichannel acquisition protocols were configured to eliminate spectral crosstalk and cross-excitation. Three-dimensional image stacks were acquired using optical zoom (3X), at 1 Airy unit pinhole, and 0.5 µm Z-interval. LSM 900 was equipped with a 63X Plan-Apochromat 1.4 NA oil immersion objective. Appropriate excitation laser lines were used for each fluorophore (as mentioned above for LSM 700). Three-dimensional image stacks were acquired using optical zoom (2X), at 1 Airy unit pinhole, and 0.5 µm Z-interval with 2X averaging. For mitochondrial potential assessment, image analysis was performed on maximum intensity projections within manually drawn Regions of interest (ROIs) around individual cells, and with appropriate background correction. The details of the newly developed mito-SinComp image analyses is described below.

### Development of mito-SinComp approach for studying structure-function relationship of single mitochondrial components

**A. Staining / expression of probes:** Modularity of the approach allows use of any appropriate probes alone or in combination. Here, we used: **a)** Structure: Mitotracker staining (as described above) for structural analyses; TMRE stain for mitochondrial potential was used while maintaining TCDD in the staining solution; **b)** Structure-function: Previously generated stable cell lines expressing validated combination of mito-PSmO2 and mito-roGFP was used for demonstrating use of mito-SinComp in live [Oxidation] function. For analyses of functional organization of mtDNA, cells or *Drosophila* tissue are co-immunostained samples Tom20 or Phb2 marking the mitochondrial structure and for DNA antibody marking mtDNA nucleoids.
**B. Microscopy:** 3D Confocal microscopy for mito-SinComp is performed (as described above) After the appropriate quality control steps, the resulting confocal micrographs were passed through mito-SinComp analytical pipeline described below.
**C. Mito-SinComp analyses:** This uses 3D confocal micrographs with appropriate combinations of compatible probes reporting mitochondrial structure and function separately. To develop mito-SinComp structure-function analysis for single mitochondrial components, we modified the open source MitoGraph code (https://github.com/vianamp/MitoGraph) to identify single mitochondrial components and include functional analyses for the identified components within single cells. Mito-SinComp analysis requires a Linux or MacOS system to run natively. It can be run on Windows through the Windows Subsystem for Linux (WSL) or another Virtual Machine application like VirtualBox. We developed mito-SinComp analysis using Ubuntu 24.04 LTS with VTK 9.0 after installing all essential libraries and packages (build-essential, cmake, git, r-base, wget, os, sys, libvtk9-dev, libgl1-mesa-dev, freeglut3-dev, qtbase5-dev, qtdeclarative5-dev, libqt5opengl5-dev, pandas, networkx, igraph, PIL, numpy (v2.1.1 for mito-SinComp and v1.1.1 for Foci Detection), tifffile, matplotlib, scikit-learn). Our customized modular script for mito-SinComp structure-function approach includes the following steps:

#### 1. Identifying the ‘structure’ and ‘function’ channels

The 3D confocal micrographs for our analysis have 3 channels, excited by 405 nm (function-1), 488 nm (function-2) and 555 nm lasers (structure). MitoGraph exclusively uses only the designated structure channel to obtain the binary image through an adaptive thresholding process^61^. Therefore, we aimed to obtain the intensity values of the ‘function’ channels in the pixels of the MitoGraph binary image. Here, we used the “splitter.py” function to separate the channels using the AICSImageIO and tifffile libraries. Each channel is then saved as an individual Z-stack in TIFF format as ‘structure’ and ‘function’ channels.

#### 2. Running MitoGraph

The open-source MitoGraph (v3.0) script is included to obtain the MitoGraph output files as described before.

#### 3. Identifying individual mitochondrial components

Mitograph produces the following three outputs containing distinct types of information-**a**) connected Components (“cc”) on interconnected elements; **b**) mitochondrial elements (“line id”) on individual mitochondrial segments; **c**) points on the mitochondrial skeleton (“point id”) on individual data points. Each “point id” belongs to a “line id”, and each “line id” belongs to a cc. This information is distributed across the following output files; 1) mitograph.txt contains information about “point id” and the corresponding “line id” they belong to; 2) mitograph.cc file links “node id” to the “cc” they belong to; **3**) mitograph.coo file provides geometric positions (x, y, z coordinates) for each “node id”. To streamline our analysis, we consolidated this data into a single, tidy data frame. To incorporate “cc” information into the mitograph.txt file for network-level analysis, we identified which “point id” entries are also “node id” entries, where nodes are the terminating points of a “line id”. Towards this, we first computed the Euclidean distance between each pair of “point id” (from mitograph.txt) and “node id” (from mitograph.coo). Thereafter, we identified the “point id” - “node id” pair with the smallest distance to identify the “point id” that corresponds to each “node id”. Once the “nodes” were identified, we used the mitograph.cc file to link the cc information to each “node id”. This enables us to append “cc” information to the mitograph.txt file. Thus, the resulting final data frame integrates “cc”, “line id”, “point id”, x, y, z coordinates and “nodes” in a single *.csv file where the structure-function features will be appended after computation as described below. Thus, a comprehensive data file is created for further downstream analyses.

#### 4. Computing structural properties for networks and non-networks

We first identified the individual networks and non-networks from the connected components (“cc”), which include both networked and non-networked components. Here, we employed a filtering approach based on the number of edges and nodes. Specifically, any connected component with one edge and two nodes, along with their corresponding data, was classified as a non-networked component. All remaining connected components, along with their corresponding data, were categorized as networked components. This classification enabled separate analysis of networked and non-networked components, facilitating a more focused examination of their distinct properties. The primary measured structural features are length and diameter, from which other features are derived. Supplementary Table-1 includes the definition and formulae of all the structural parameters studied.

#### 5. Computing redox function for networks and non-networks in live cells

The redox function was computed by obtaining the average pixel intensity ratio of individual cc by mapping the ‘function’ information to the ‘structure’ information obtained as above. Thereafter, the redox functional property of individual mitochondrial component was computed and finally validated (**Fig. 2C; Fig. S1G-H**). The details of these steps are below:

##### i. Converting MitoGraph output to pixel coordinates

MitoGraph was run using Z-stack for the structure channel using runMitograph.py function in a batch processing mode for all cells within a single directory. MitoGraph outputs the geometric coordinates (x, y, z) of the mitochondrial skeleton as per the provided scale when running MitoGraph. The pixel coordinates (x_pixel, y_pixel, z_pixel) can be retrieved from the geometric coordinates through the scale using the formula, where D represents either x, y, or z. This is then rounded to the nearest integer, as pixel coordinates are a discrete quantity.

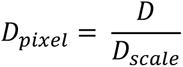

**Fig. S9A** shows an example of the results of this process, where the scale for x and y is 0.104 and is 0.5 for z. The conversion is confirmed by the overall overlap of the computed pixel coordinates (in red) with the MIP of the structural channel (**Fig. S9B**).

##### ii. Computing the intensity values of the pixel coordinates from individual ‘function’ channels

The Z-stacks Tiff images are converted into a three-dimensional array using tifffile library. The intensity value of each pixel coordinate generated in step-i is computed from the individual functional channels. The average intensity of the 9 pixels in a 1X3X3 voxel is computed and assigned to the central pixel. This is repeated for every x, y, z pixel computed in step-i. In the following example, in the slice of an image at z = 2, the intensity of a_222_ is the average of all the colored pixels.

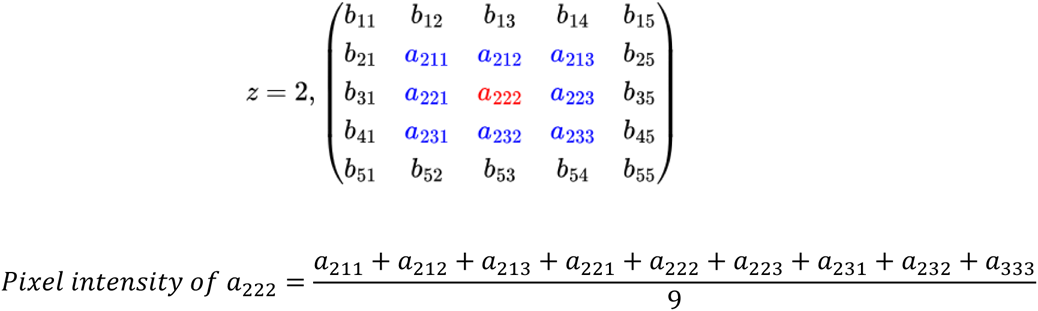

##### iii. Computing pixel intensity ratio to reflect redox function

Once the intensity for each pixel is obtained (step ii), it is put into the “pixel_intensity_channel_N” column, where “N” reflects the selected channel. For computing pixel intensity for individual element, the intensities for each channel are grouped according to their “line_id” and averaged. These average intensity values were used to compute pixel intensity ratio at the element level. For compute the pixel intensity ratio at the individual cc level (networks or non-networks), the pixel intensity ratio of the elements comprising the cc are grouped according to their “cc_id” and averaged to find the pixel intensity ratio of the “cc”, as exemplified for an individual non-networked “cc” (see **Fig.S9C** and equations below). These values are validated (see main text) using appropriate modules in python (scikit-learn).

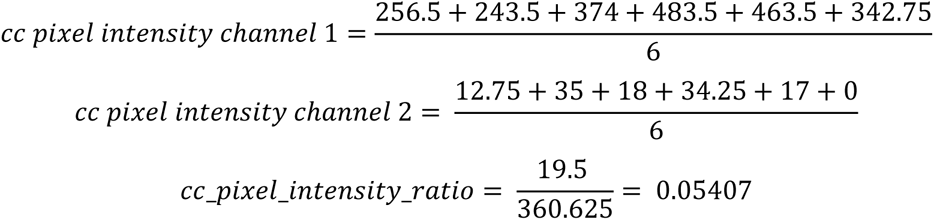

##### iv. Visualizing single networks and non-networks within single cells in 3D graphs

Visualization of the data from the comprehensive data file is performed using the 3D_generate_graphs.py script. The data is plotted by looping through each “cc” and each “line id”. The individual points in each “line id” are then plotted to ensure that the order of the points is preserved in the visualization. To distinguish between shapes, each “cc” is assigned a random and unique color. These shapes are then rendered in a 3D space. The same process is repeated for generating the LUT of the ‘function’ values, where the color code represents the [Oxidation] function of individual elements.

#### 6. Computing mtDNA related functional features for networks and non-networks in fixed samples

False positive signal from the nucleus was excluded by manually drawing a region of interest (ROI) around the nucleus using ImageJ. Such processed 3D confocal micrographs were analysed using our Python-based pipeline created including DBSCAN (Density-Based Spatial Clustering of Applications with Noise) algorithm for detecting the mtDNA signal mapped or unmapped to the mitochondrial marker. The following steps are executed after importing all necessary packages (mentioned in Step C of Methods):

##### i. Intensity based thresholding and Density-based clustering

The structural and functional channels are loaded as separate tiff files. Image J is used to define the threshold from the MIP of the functional channel to include both bright and dim mtDNA nucleoids. Next, the image is converted to binary where mtDNA signal appeared as foci. The 3D coordinates of the foci are obtained using DBSCAN algorithm. For DBSCAN clustering, two defined parameters are considered: epsilon (ε) - defines the maximum distance between two points to be in one cluster and minimum sample - minimum number of pixels above the threshold in a cluster. In our study, ε was set to 1 and the minimum sample to 4 by visual comparison of the DBSCAN output coverage with the mtDNA distribution in the raw image.

##### ii. Defining quality check of clusters

A unique list of clusters that contain one point is created and their coordinates are obtained. Thereafter, a 3×3 square is drawn on gray scale image using each coordinate as the center, and the mean pixel intensity of that square is computed. If the mean pixel intensity of that square is below 90% of the pixel intensity of the center, the coordinate is removed. The coordinates in each cluster are obtained and stored in the form of a pandas data frame.

##### iii. Extracting cluster information

The clusters are renamed with Z slice number as prefix in *.csv file with filtered coordinates. The cluster size is then computed as the number of pixels in one cluster and saved the updated data frame to a new *.csv file. To obtain cluster coordinate, size and number the data of the *.csv is loaded, ensuring that the necessary columns are in the data frame. To remove the pixels labelled as noise (−1) during the quality check were filtered out from the slice of interest. This data is then saved in a separate *.csv file and turned into the final data frame to be considered for mapping.

##### iv. Mapping of functional coordinates to the structural coordinates

This involves loading and extracting coordinates from the function channel data frame created in the previous step and the mito-SinComp structure channel data frame, separately. Next, the pairwise Euclidean distances of each coordinate of the above data frames are computed and the minimum distance is considered as ‘mapped’. In this study, we optimized this minimum distance as ≤10 by visually comparing DBSCAN output coverage with the mtDNA distribution in the raw image. Therefore, if any DBSCAN cluster is beyond 6 pixels of the structural component was considered unmapped, and the rest mapped. The final functional channel data frame for both mapped and unmapped clusters was saved as *.csv separately. Finally, the mapped and unmapped clusters obtained from individual z-slice are visualized by matching the spatial coordinate information of the clusters (function) separately onto the structural channel MIP.

##### v. Computing and storing cluster level information from a single slice with structural level information

The structure channel data frame and the last updated functional channel data frame are merged on index followed by computing functional metrics like cluster number, cluster size, and average cluster size per cc_x. This merged data frame is saved as final *.csv file for further downstream analysis.

#### 7. Data visualization and representation

The output of mito-SinComp analyses is a comprehensive data file (*.csv) holding the relevant details for individual mitochondria in individual cells; rows with point_ids under line_ids under cc_ids while columns with relevant values for each id. This file is used to obtain and visualize various structural-function parameters. For inter- and intracellular comparisons of SMNs and HMNs, the *.vtk files generated by MitoGraph for individual cells were analyzed and visualized using the open-source software ParaView-v5.13.0. All plots were generated in Python using appropriate modules.

### Identification of novel structure metric-[CRL mito-Net] to distinguish vSMNs, SMNs and HMNs

We employed a thresholding approach based on the [CRL mito-Net] metric in combination with the previously defined cellular [Fusion1] metric, which represents the percentage of the longest component length within a cell. The cell with the minimum Fusion1 value was identified in the HF-cells group. Within this cell, the network with the highest [CRL mito-Net] value was used as a threshold (−1.4) to demarcate the boundary between the SMN and HMN zones. To define the SMN region, we visually assessed the peak of this metric within our positive control INT-Cell group, which showed a significant enrichment of SMNs. This assessment identified a threshold of - 4.1, with any values equal to or below this point categorized as vSMNs. This approach effectively divides the entire distribution into three distinct zones based on the [CRL mito-Net] metric. The same thresholding criteria were consistently applied across different cell lines and experimental conditions to study mitochondrial networks, with a primary focus on SMNs and HMNs. Given the versatility and robustness of this dimensionless metric, it can be adapted for thresholding in future studies, depending on the specific research question or the mitochondrial components of interest, including both networked and non-networked mitochondrial structures.

### Radom Forest based prediction of mitochondrial function from structure

The random forest machine learning algorithm is an ensemble learning method that performs regression by constructing multiple decision trees during training. Each tree is trained on a bootstrapped subset of the data, and node splits are determined using a random subset of the features in the dataset. The final prediction is obtained by averaging the predictions of all individual trees.

We implemented the random forest model using the ‘scikit-learn’ package (version 1.5.2). The key parameters optimized during training included: ‘n_estimators’: The number of trees in the forest; ‘max_depth’: The maximum depth of each tree; ‘ccp_alphà: The complexity parameter used for minimal cost-complexity pruning. The algorithm selects features and split points based on the reduction in variance for regression tasks. While training the model, 70% of the data was used as the training set and 30% was used as the testing set. A random seed was set to ensure all experiments were reproducible. Default values were used for parameters not explicitly mentioned unless otherwise stated.

We performed hyperparameter tuning using the ‘BayesSearchCV’ function from the ‘scikit-optimizè (‘skopt’) library. This method implements Bayesian optimization to efficiently explore the hyperparameter space, sampling a fixed number of parameter combinations rather than exhaustively evaluating all possibilities as in grid search. The search was configured with the following parameters: Search space: Defined by specifying distributions for each hyperparameter (‘n_estimators’, ‘max_depth’, ‘ccp_alphà); n_iter: The number of parameter settings sampled from the distributions. We used a value of 100. Default values were used for unspecified parameters unless otherwise noted.

We assessed model performance using the coefficient of determination (R²) on the held-out test set. R² is computed as:

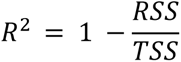

Where RSS is the sum of squares of residuals and TSS is the total sum of squares. R² quantifies the proportion of variance in the target variable explained by the model, with values closer to 1 indicating better fit.

We evaluated model performance using 5-fold cross-validation, implemented via the ‘cross_val_scorè function from ‘scikit-learn’. The training set was partitioned into five equal-sized folds, each preserving the overall distribution of the target variable. The model was trained on four folds and validated on the remaining fold, with this process repeated five times such that each fold served as the validation set once. The R² score were computed for each validation fold and aggregated to produce an average score with standard deviation. This approach provided an estimate of generalization error while mitigating bias from any single train-test split. The same cross-validation folds were used consistently during hyperparameter optimization and final evaluation to ensure comparability.

To interpret the model, we analyzed feature importance scores from the trained random forest using the ‘feature_importances_’ attribute of the Random Forest Regressor class. This attribute computes scores as the mean decrease in impurity (Gini importance) across all trees, reflecting each feature’s relative contribution to predictive accuracy. The importance values are normalized to sum to 1 for comparative analysis. Performance metrics and feature importance were computed using native ‘scikit-learn’ implementations to ensure consistency with the training process. The test set was not used during hyperparameter optimization or feature selection to prevent data leakage.

### High Dimensional weighted gene correlation network analyses (hdWGCNA)

#### Single-Cell RNA Seq

TCDD exposed cells were processed for scRNA-seq using the 10X Genomics 3’ v3.1 NextGem platform and analyses were performed using Seurat (v4.3.3). Notably, the cells were tested for > 90% viability before scRNA-seq. Libraries were constructed through cDNA fragmentation, end repair, adaptor ligation, and dual-indexed PCR amplification. Sequencing was performed on Illumina Nextseq500 (28 bp R1, 90 bp R2, 10 bp i7/i5 indices) with ≥20,000 reads/cell. Only cells expressing a minimum of 2000 detectable genes were included using the ‘nFeature_RNA’ filter and variable feature sets were identified by *FindVariableFeatures* function in Seurat. This filtering also excluded dead cells with >10% mtDNA gene expression in all our datasets analyzed. This allowed us to exclude the percent.mt filter that uses mtDNA-based exclusion of dead cells. *nFeature_RNA* >2000 was applied prior to normalization of data using the *NormalizeData* function in Seurat. Approximately ∼3000 genes in an average of 5000 cells were covered in our data in each population. The same filters were used for processing publicly available raw data from cSCC patients^50^. Specific sets of samples were combined using the *IntegrateData* function for the outlined comparison sets - transformed, exposed, DRP1 knockdown and cSCC resulting in four sets of scRNASeq analyses. These integrated datasets were scaled and clustering of cells was performed with up to 20 dimensions (dim = 1:20) with 0.2 resolution quantified for stem cell high cluster identification, visualized by UMAP. Marker genes for each cluster were identifed with *FindAllMarkers* function with statistical significance calculation performed using by Wilcoxon Rank Sum test. While, the FindMarkers function (pseudobulk analyses) of Seurat was used to compute the differential expression of the cluster markers (normalized data before integration (assay ‘RNA’)) using the Wilcoxon rank sum test. Average Expression was computed for select genes of interest using the *Average Expression* function. GSEA analyses were performed for scRNASeq clusters with ranked log fold change (LFC) of markers (LFC > 0.1, p < 0.01, FDR q value p < 0.01) using MsigDB and Reactome databases. To sum up, Seurat objects from the resultant count matrices were processed, integrated, visualized, and analyzed for relevant downstream analyses including pairwise differential gene expression analyses (using FindMarkers function), cell cluster analyses (using FindAllMarkers function), and Gene Set Enrichment Analysis (GSEA) for cell cluster annotation. Immunoblotting was performed to validate key scRNAseq finding using standard methods using commercial antibodies (Key Resource Table).

#### hdWGCNA design

We applied method on scRNA-seq data to identify transcriptomic networks^62,63^ following the workflow outlined in **Fig. S3B**. We used integrated Seurat objects for both pseudobulk and cell cluster-based analyses, considered the top 20 / 25 genes (ranked by kME) to visualize transcriptomic interactions through hub-gene network plots. Functional annotation of the hdWGCNA modules was performed using GSEA where the modules with ≥ 5 mtDNA genes as the leading-edge genes were reassigned as ‘mtDNA’ in our analyses. Edge distance and the number of edges were quantified using TOM, providing insights into gene connectivity and interaction strength.

#### Detailed workflow of hdWGCNA

hdWGCNA^62^(smorabit.github.io/hdWGCNA/) was used to identify gene interaction modules amongst differentially expressed genes (> 0.25 avg_log2FC, <10^-5^ p_val_adj, pct ≥ 0.25 in pseudobulk analyses using FindMarkers function) and amongst the characteristic marker genes for the identified cell clusters (> 0.25 avg_log2FC, <10^-5^ p_val_adj). QC plots of nCounts_RNA distributions were used to establish lower thresholds for outlier removal (**Fig. S9D).** This processed integrated Seurat object is used as an input for respective hdWGCNA, both for pseudobulk and cell cluster levels. The hdWGCNA object was initialized, and metacells were constructed using K-nearest neighbors (KNN) (**Figs. S9E-F**). Gene expression matrices were constructed, and pairwise Pearson’s correlations were computed, transformed into adjacency matrices using a soft-thresholding power approach (**Fig. S9G**), and used to construct networks using Topological Overlap Measures (TOM) values. The hdWGCNA algorithm ensured a scale-free topology during signed network construction for each hdWGCNA object for both pseudobulk and cell cluster. Transcriptomic modules were detected using hierarchical clustering and TOM, with module eigengenes (MEs) computed for further analysis. Module connectivity (kME) values were computed to identify hub genes characterized by high kME values. Selection of specific parameter cut-offs for hdWGCNA are elaborated below: **a.** K-nearest neighbors (KNN)-based dimensionality reduction: Metacells for each experimental/control group (pseudobulk) and specific clusters were constructed using KNN, leveraging biological similarity to define cellular subpopulations in each hdWGCNA object.

Pearson correlation-based density plots guided the selection of K value of 20 with a maximum of 10 shared neighbors for an average of 5000 cells in each population, optimizing the balance between capturing local structure and minimizing noise in sparse data (**Figs. S9E-F**); **b.** Soft Thresholding Power (β): The optimal β value was selected by evaluating the scale-free topology fit index, to enhance strong correlations and reduce weaker ones, aiming for the lowest power where the network closely approximated a scale-free structure. The β values between 6 and 10 were determined as optimal, balancing network sparsity with scale-free topology at both pseudobulk and cell cluster-based hdWGCNA objects (solid black circles on plots in (**Fig. S9G**). This transformation enabled robust, biologically meaningful signed networks across cellular subpopulations; **c.** Module Connectivity (kME): kME measures the correlation between a gene’s expression profile and its module eigengene, indicating the gene’s alignment with the module’s expression pattern. High kME values suggest that a gene has a central role within the module, while low values indicate a less integral role. Genes in each module were ranked by kME, identifying the top 20 in pseudobulk (**Fig. 3E, S3D, S6B-C**) and top 25 (**Fig. 5F, 5K, S7D**) in cell cluster-based hdWGCNA for functional significance. This kME analysis highlighted closely interacting clusters, enabling us to identify top-ranking hub genes within a network.

### Statistical Analyses

Individual experiments were performed at least 3 times, and appropriate statistics were applied. The N detected for individual mitochondrial component per cell ranges from 30 to >100. N for each analysis is mentioned in the figure and can be scaled as necessary. Appropriate statistical analyses for single cell microscopy were conducted using either Excel or Python (SciPy); The Kolmogorov-Smirnov test was applied to histogram and ridge plots; the Mann-Whitney U test to violin and box plots; the Student’s t-Test to bar plots; and the Chi-square test to bivariate scatter plots. Corresponding p-values were computed and adjusted for multiple hypotheses testing using the Holm-Bonferroni correction method. Bootstrapping was used for resampling, as needed. The hdWGCNA and Random Forest used inbuilt statistical analyses as required.

## Data Availability

The raw, analyzed and meta data of scRNA-seq experiment have been deposited in the Gene expression Omnibus (accession number GSE285728). All data matrices, scripts, and hdWGCNA results will be deposited in public repositories upon publication, to ensure transparency and reproducibility for future research.

## Acknowledgements

The research, D.P. and B.S were supported by NIH (R33 ES025662/ES/NIEHS) and DBT-Wellcome Trust India Alliance (IA/S/20/2/505198/WTDBT); S.A. was supported by DBTRA/2023-24/NE/Ashoka/19; M.S. was supported by DBT-JRF-DBT/2023-24/Ashoka Uni/2196; B.W. was supported by (IA/S/20/2/505198/WTDBT); B.G. was supported by DBT/2022-23/ASHOKA/2156; PS was supported by Mphasis F1 Foundation. We acknowledge UAB Flow Cytometry and Single Cell Services Core (supported by P30 AR048311, P30 AI027667) and Dr. Mike Crowley from the Heflin Genomics Core for scRNA-seq, we sincerely thank Dr. LS Shashidhara, Dr. Alok Bhattacharya and Dr. Gautam Menon for critically reading the manuscript, and Dr. KM Poonam for crosschecking RF code set.

## Key Resources Table

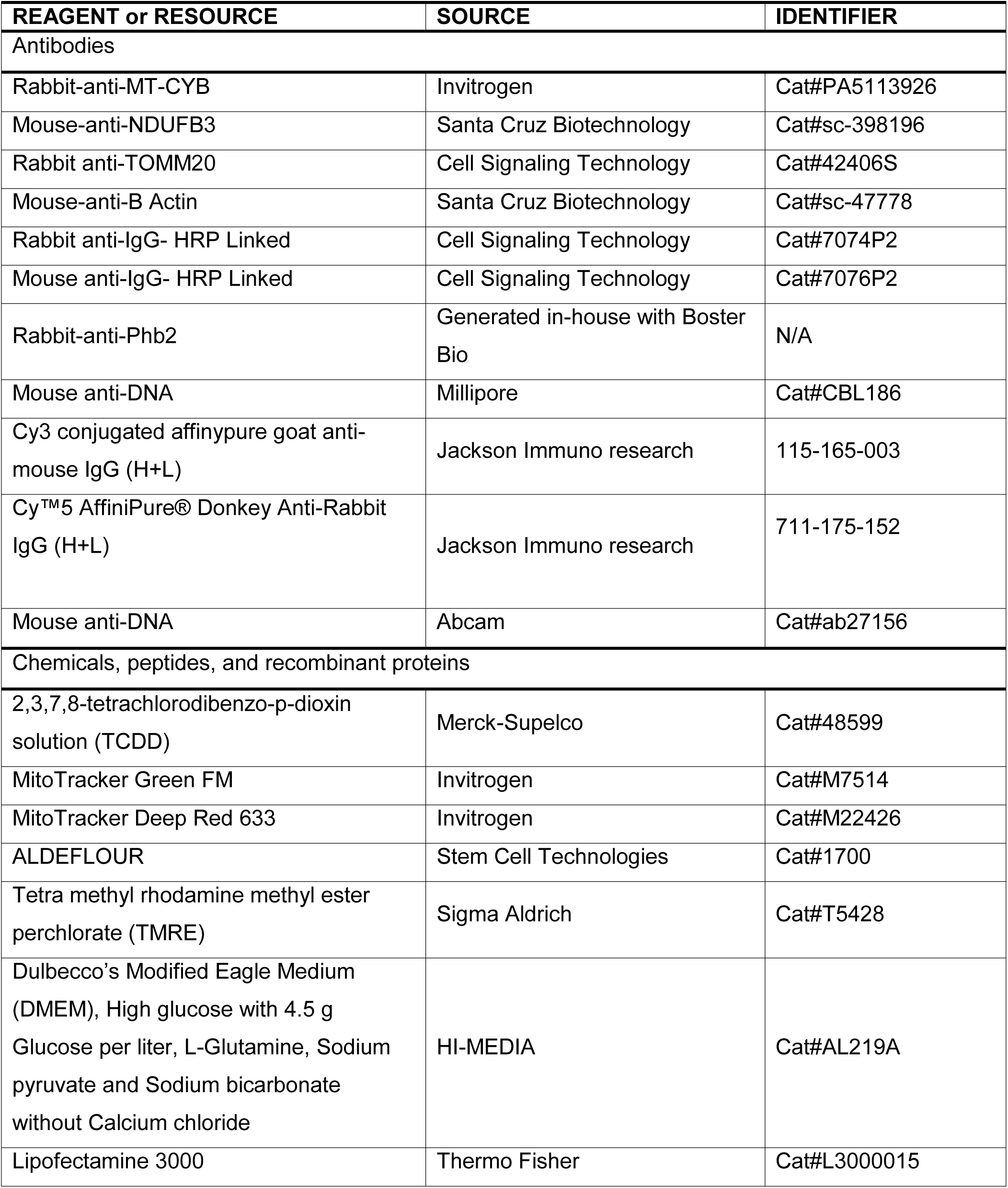
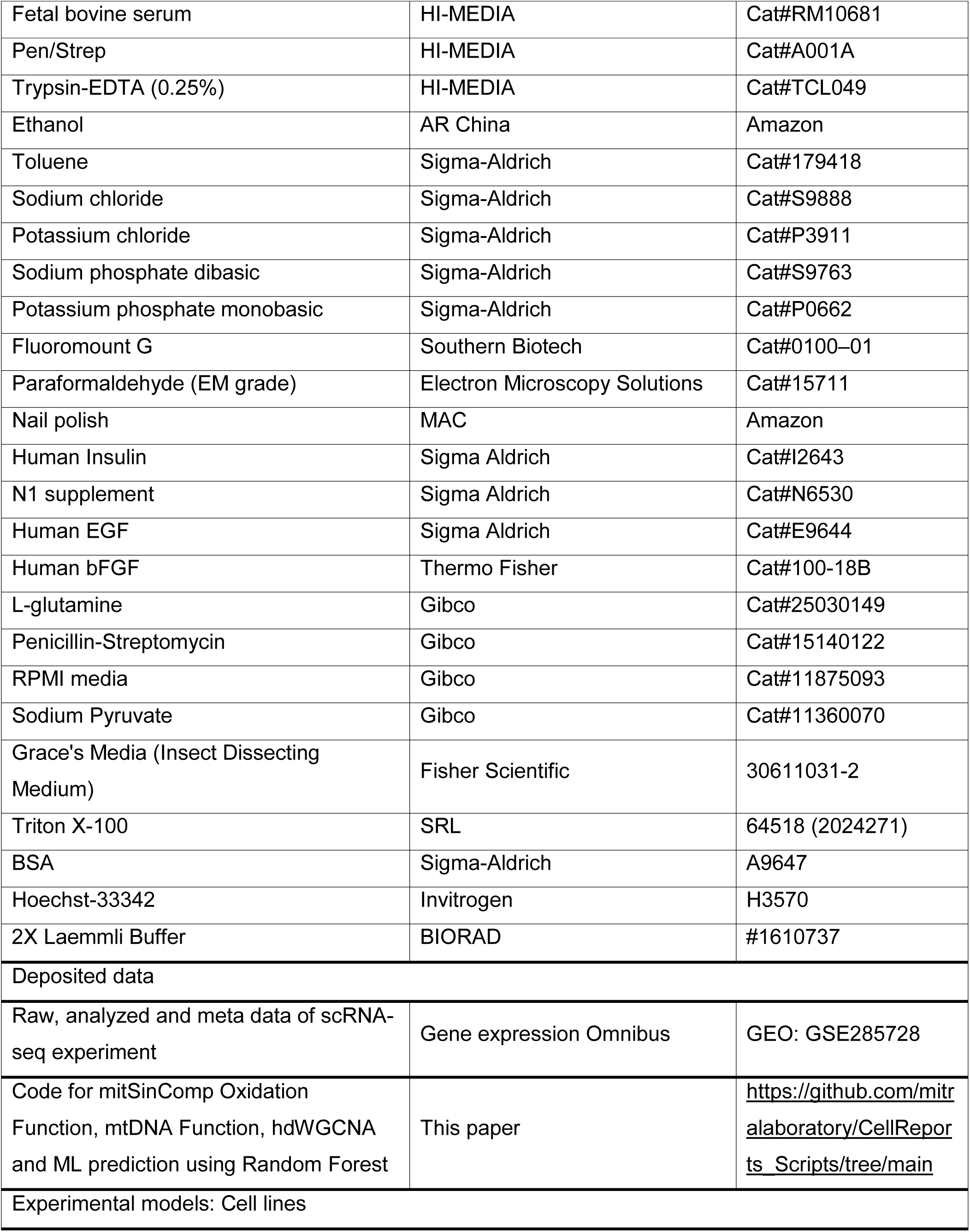
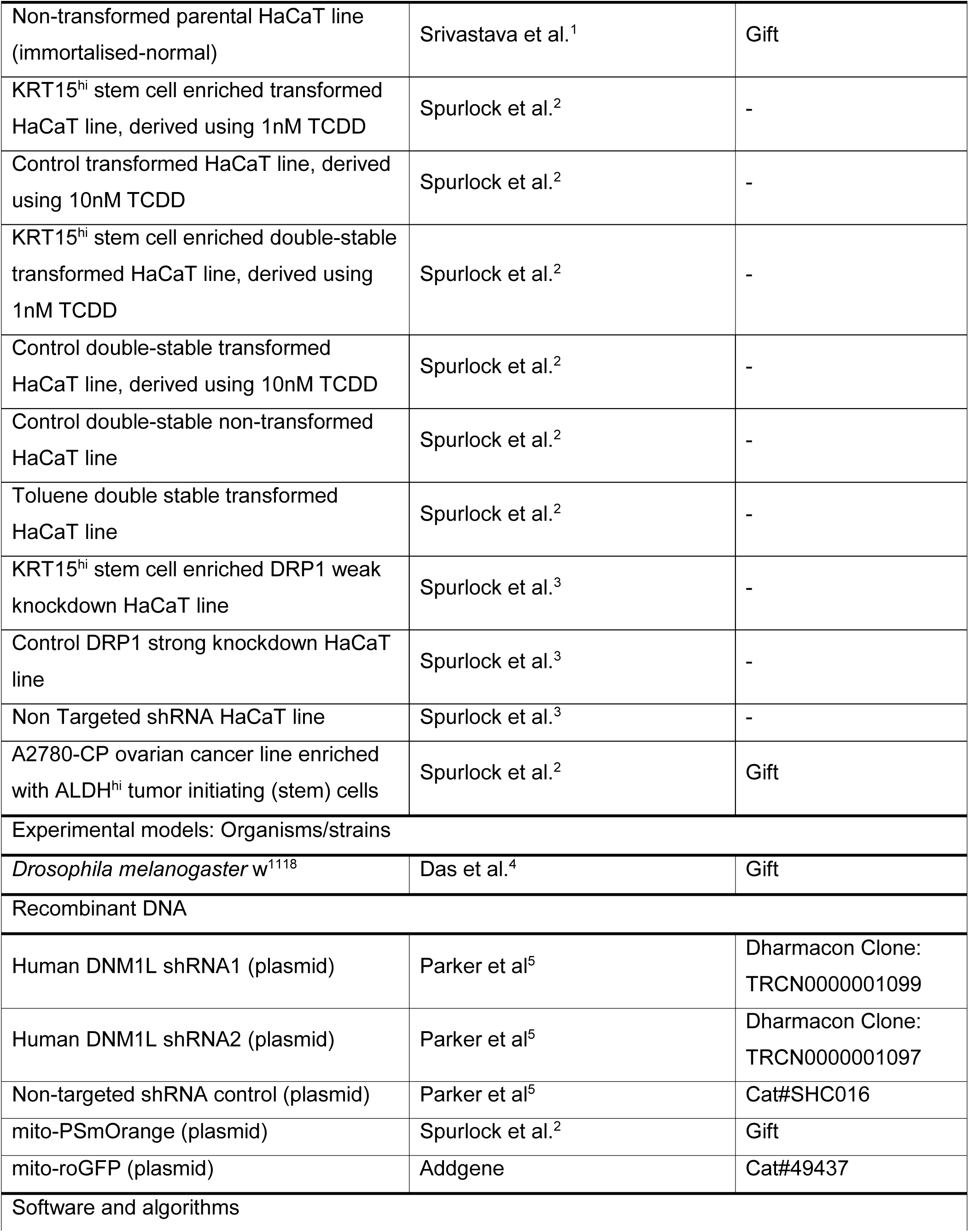
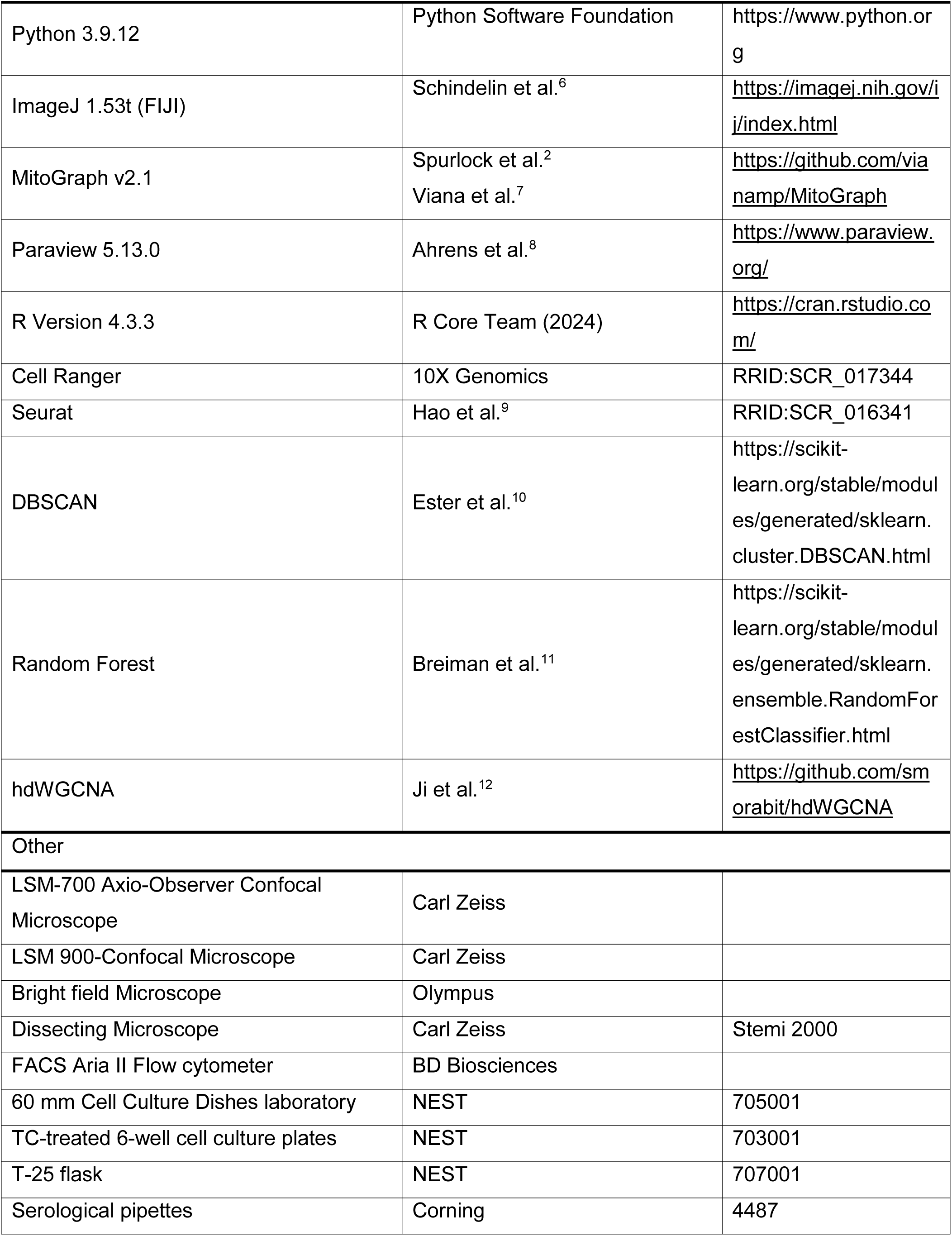
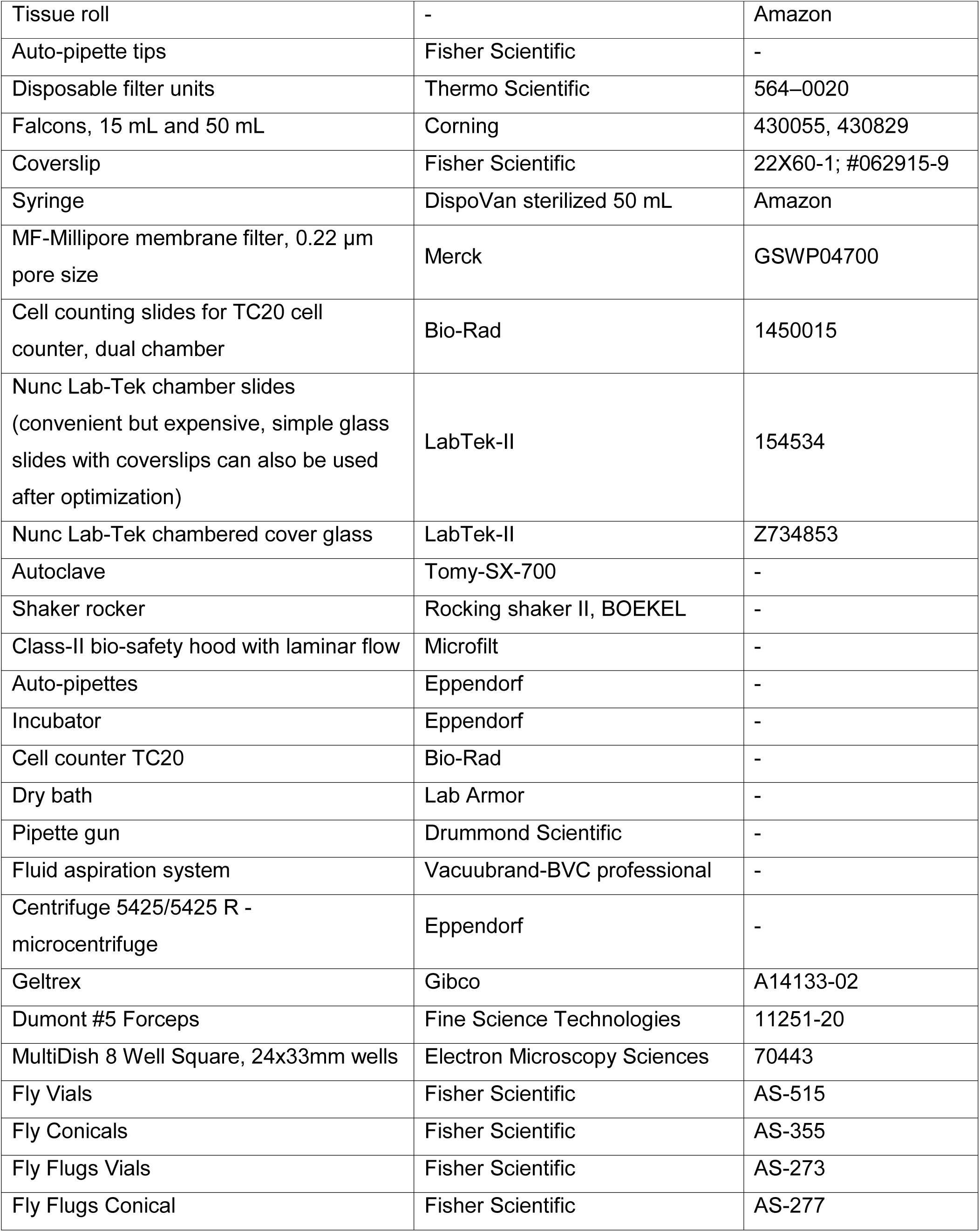

