## Supplementary material for "Quantitative analyses of single mitochondrial structure-function heterogeneity uncovers stemness specification by redox-tuned small-mitochondrial-networks": Supplemntal figures and legends

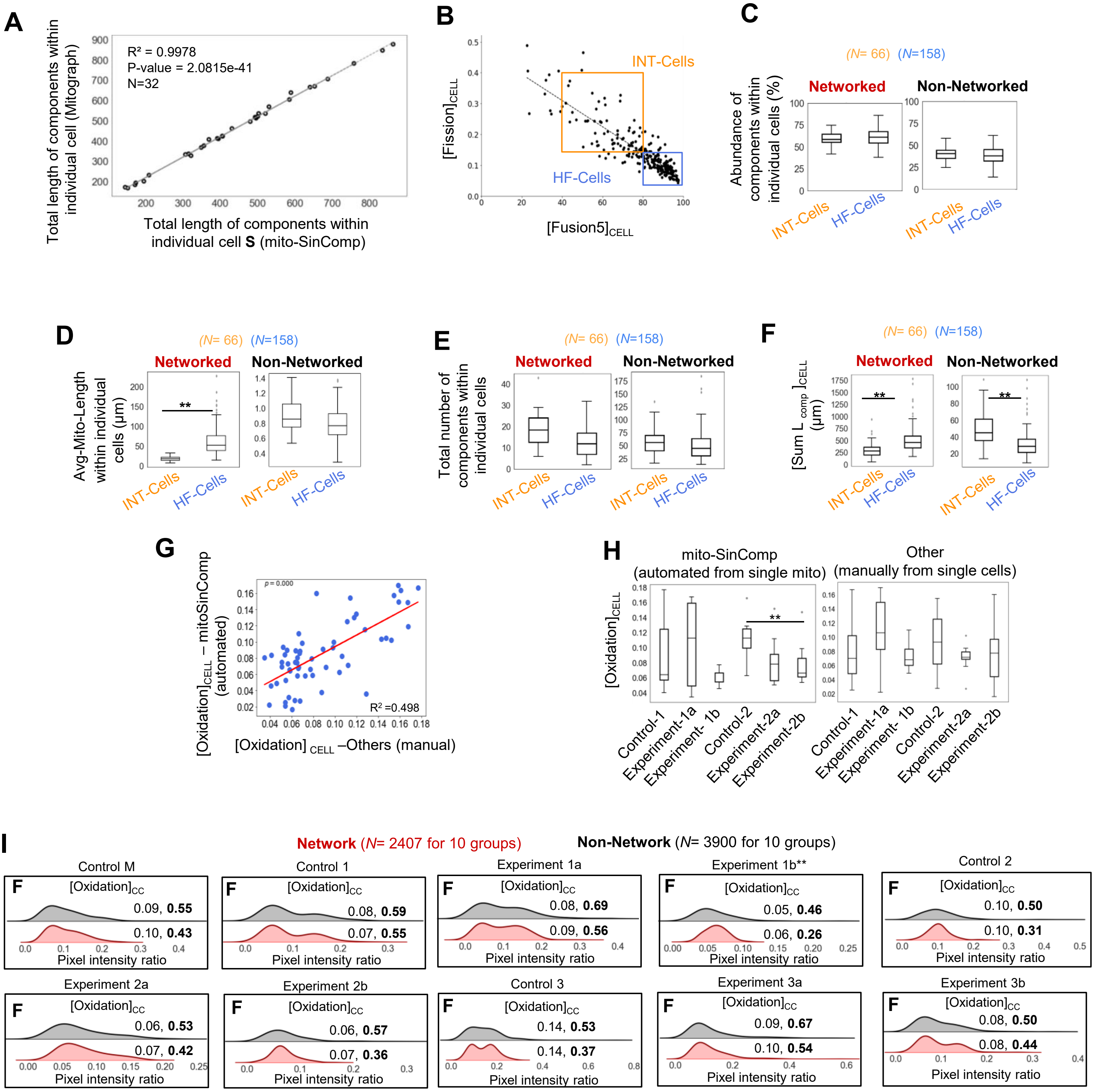
A
R² = 0.9978
P-value = 2.0815e-41
N=32
Total length of components within individual cell (Mitograph)
Total length of components within individual cell S (mito-SinComp)
B
INT-Cells
HF-Cells
[Fission]CELL
[Fusion5]CELL
C
(N= 66) (N=158)
Networked
Non-Networked
Abundance of components within individual cells (%)
INT-Cells
HF-Cells
D
(N= 66) (N=158)
Networked
Non-Networked
Avg-Mito-Length within individual cells (µm)
**
INT-Cells
HF-Cells
E
(N= 66) (N=158)
Networked
Non-Networked
Total number of components within individual cells
INT-Cells
HF-Cells
F
(N= 66) (N=158)
Networked
Non-Networked
[Sum L comp ]CELL (µm)
**
INT-Cells
HF-Cells
G
p = 0.000
R² =0.498
[Oxidation]CELL – mitoSinComp (automated)
[Oxidation] CELL –Others (manual)
H
mito-SinComp
(automated from single mito)
Other
(manually from single cells)
[Oxidation]CELL
**
Control-1
Experiment-1a
Experiment- 1b
Control-2
Experiment-2a
Experiment-2b
I
Network (N= 2407 for 10 groups)
Non-Network (N= 3900 for 10 groups)
Control M
[Oxidation]CC
0.09, 0.55
0.10, 0.43
Pixel intensity ratio
Control 1
0.08, 0.59
0.07, 0.55
Experiment 1a
0.08, 0.69
0.09, 0.56
Experiment 1b**
0.05, 0.46
0.06, 0.26
Control 2
0.10, 0.50
0.10, 0.31
Experiment 2a
0.06, 0.53
0.07, 0.42
Experiment 2b
0.06, 0.57
0.07, 0.36
Control 3
0.14, 0.53
0.14, 0.37
Experiment 3a
0.09, 0.67
0.10, 0.54
Experiment 3b
0.08, 0.50
0.08, 0.44

Fig. S1

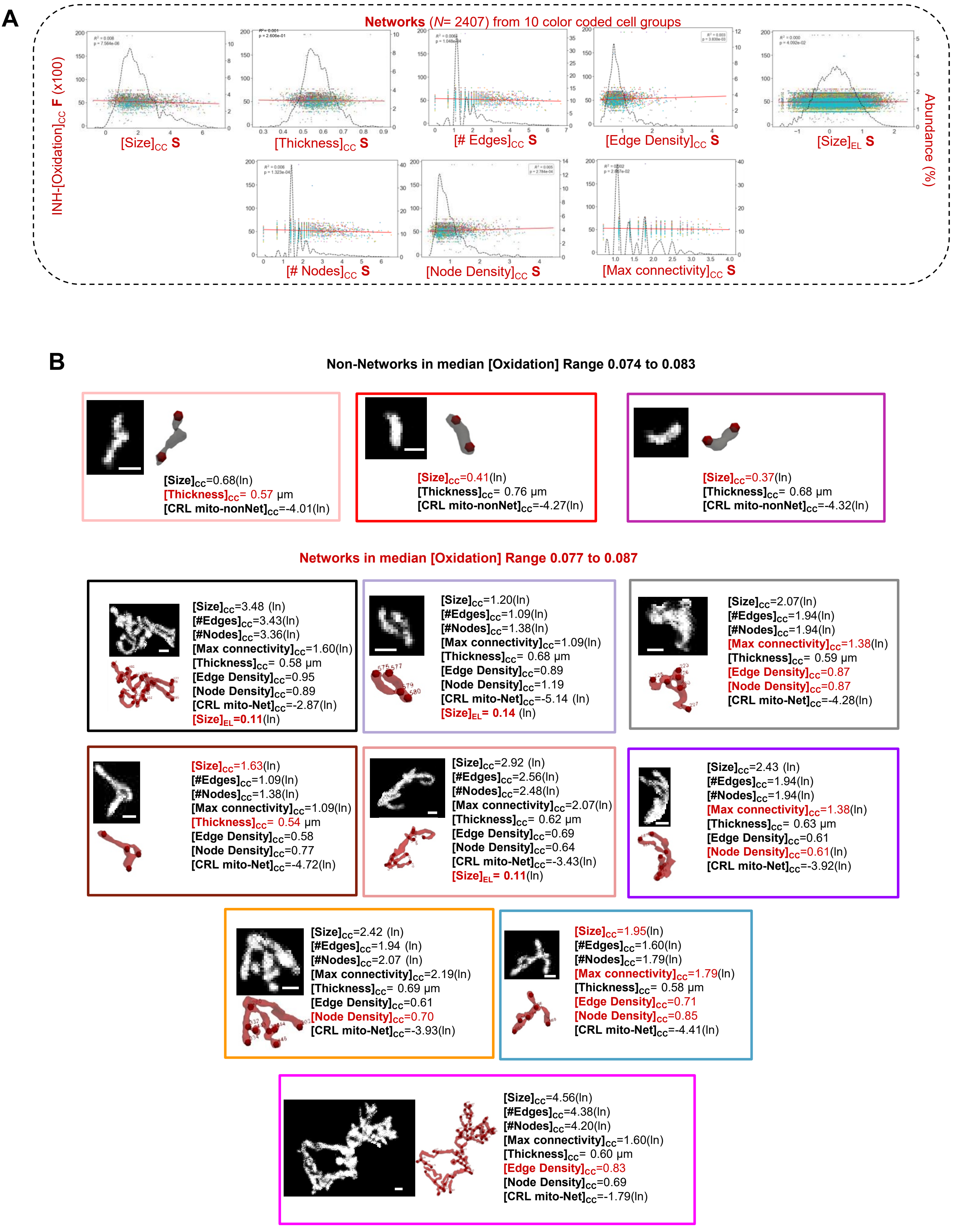
A
Networks ($N$= 2407) from 10 color coded cell groups
INH-[Oxidation]$_{CC}$ F (x100)
Abundance (%)
[Size]$_{CC}$ S
[Thickness]$_{CC}$ S
[# Edges]$_{CC}$ S
[Edge Density]$_{CC}$ S
[Size]$_{EL}$ S
[# Nodes]$_{CC}$ S
[Node Density]$_{CC}$ S
[Max connectivity]$_{CC}$ S
B
Non-Networks in median [Oxidation] Range 0.074 to 0.083
[Size]$_{CC}$=0.68(ln)
[Thickness]$_{CC}$= 0.57 μm
[CRL mito-nonNet]$_{CC}$=-4.01(ln)
[Size]$_{CC}$=0.41(ln)
[Thickness]$_{CC}$= 0.76 μm
[CRL mito-nonNet]$_{CC}$=-4.27(ln)
[Size]$_{CC}$=0.37(ln)
[Thickness]$_{CC}$= 0.68 μm
[CRL mito-nonNet]$_{CC}$=-4.32(ln)
Networks in median [Oxidation] Range 0.077 to 0.087
[Size]$_{CC}$=3.48 (ln)
[#Edges]$_{CC}$=3.43(ln)
[#Nodes]$_{CC}$=3.36(ln)
[Max connectivity]$_{CC}$=1.60(ln)
[Thickness]$_{CC}$= 0.58 μm
[Edge Density]$_{CC}$=0.95
[Node Density]$_{CC}$=0.89
[CRL mito-Net]$_{CC}$=-2.87(ln)
[Size]$_{EL}$=0.11(ln)
[Size]$_{CC}$=1.20(ln)
[#Edges]$_{CC}$=1.09(ln)
[#Nodes]$_{CC}$=1.38(ln)
[Max connectivity]$_{CC}$=1.09(ln)
[Thickness]$_{CC}$= 0.68 μm
[Edge Density]$_{CC}$=0.89
[Node Density]$_{CC}$=1.19
[CRL mito-Net]$_{CC}$=-5.14 (ln)
[Size]$_{EL}$= 0.14 (ln)
[Size]$_{CC}$=2.07(ln)
[#Edges]$_{CC}$=1.94(ln)
[#Nodes]$_{CC}$=1.94(ln)
[Max connectivity]$_{CC}$=1.38(ln)
[Thickness]$_{CC}$= 0.59 μm
[Edge Density]$_{CC}$=0.87
[Node Density]$_{CC}$=0.87
[CRL mito-Net]$_{CC}$=-4.28(ln)
[Size]$_{CC}$=1.63(ln)
[#Edges]$_{CC}$=1.09(ln)
[#Nodes]$_{CC}$=1.38(ln)
[Max connectivity]$_{CC}$=1.09(ln)
[Thickness]$_{CC}$= 0.54 μm
[Edge Density]$_{CC}$=0.58
[Node Density]$_{CC}$=0.77
[CRL mito-Net]$_{CC}$=-4.72(ln)
[Size]$_{CC}$=2.92 (ln)
[#Edges]$_{CC}$=2.56(ln)
[#Nodes]$_{CC}$=2.48(ln)
[Max connectivity]$_{CC}$=2.07(ln)
[Thickness]$_{CC}$= 0.62 μm
[Edge Density]$_{CC}$=0.69
[Node Density]$_{CC}$=0.64
[CRL mito-Net]$_{CC}$=-3.43(ln)
[Size]$_{EL}$= 0.11(ln)
[Size]$_{CC}$=2.43 (ln)
[#Edges]$_{CC}$=1.94(ln)
[#Nodes]$_{CC}$=1.94(ln)
[Max connectivity]$_{CC}$=1.38(ln)
[Thickness]$_{CC}$= 0.63 μm
[Edge Density]$_{CC}$=0.61
[Node Density]$_{CC}$=0.61(ln)
[CRL mito-Net]$_{CC}$=-3.92(ln)
[Size]$_{CC}$=2.42 (ln)
[#Edges]$_{CC}$=1.94 (ln)
[#Nodes]$_{CC}$=2.07 (ln)
[Max connectivity]$_{CC}$=2.19(ln)
[Thickness]$_{CC}$= 0.69 μm
[Edge Density]$_{CC}$=0.61
[Node Density]$_{CC}$=0.70
[CRL mito-Net]$_{CC}$=-3.93(ln)
[Size]$_{CC}$=1.95(ln)
[#Edges]$_{CC}$=1.60(ln)
[#Nodes]$_{CC}$=1.79(ln)
[Max connectivity]$_{CC}$=1.79(ln)
[Thickness]$_{CC}$= 0.58 μm
[Edge Density]$_{CC}$=0.71
[Node Density]$_{CC}$=0.85
[CRL mito-Net]$_{CC}$=-4.41(ln)
[Size]$_{CC}$=4.56(ln)
[#Edges]$_{CC}$=4.38(ln)
[#Nodes]$_{CC}$=4.20(ln)
[Max connectivity]$_{CC}$=1.60(ln)
[Thickness]$_{CC}$= 0.60 μm
[Edge Density]$_{CC}$=0.83
[Node Density]$_{CC}$=0.69
[CRL mito-Net]$_{CC}$=-1.79(ln)

**Fig. S2**

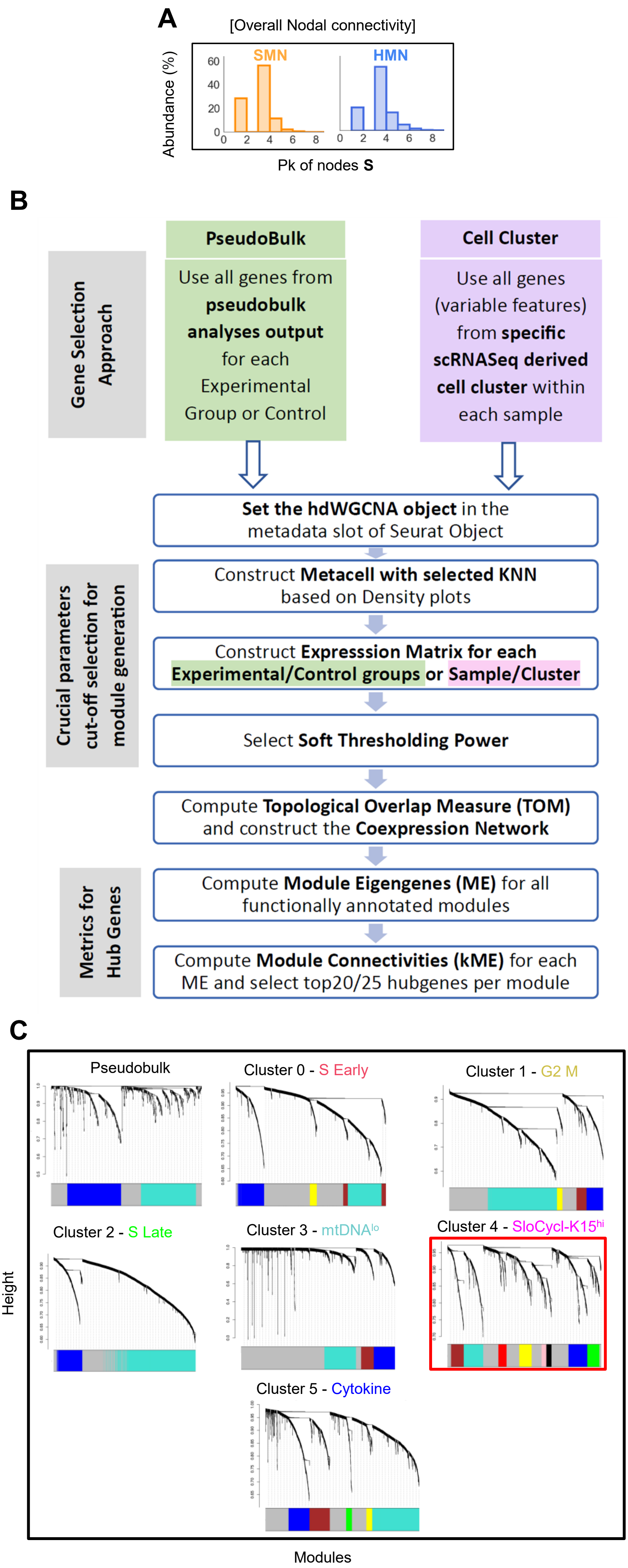


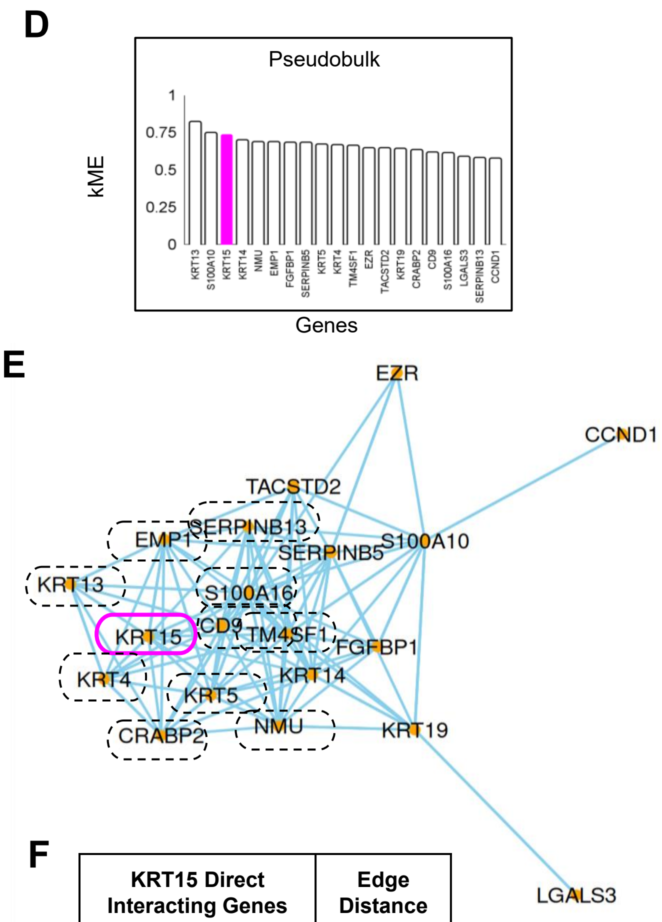


F

| KRT15 Direct Interacting Genes | Edge Distance |
|---|---|
| CD9 | 0.93 |
| KRT5 | 0.92 |
| NMU | 0.66 |
| CRABP2 | 0.53 |
| TM4SF1 | 0.50 |
| KRT13 | 0.44 |
| S100A16 | 0.39 |
| KRT4 | 0.35 |
| EMP1 | 0.29 |
| SERPINB13 | 0.28 |

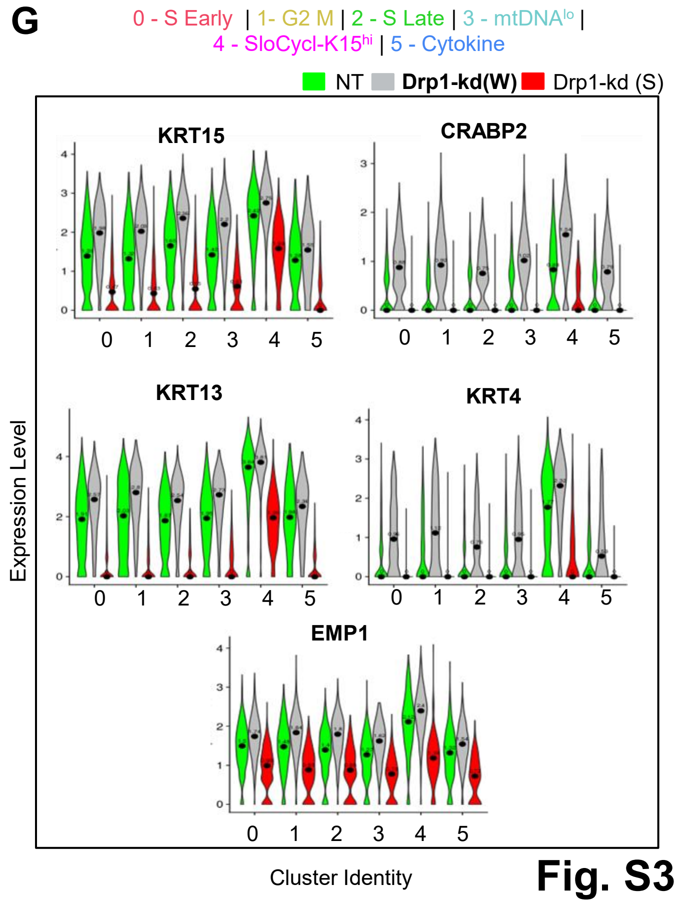


**Fig. S3**

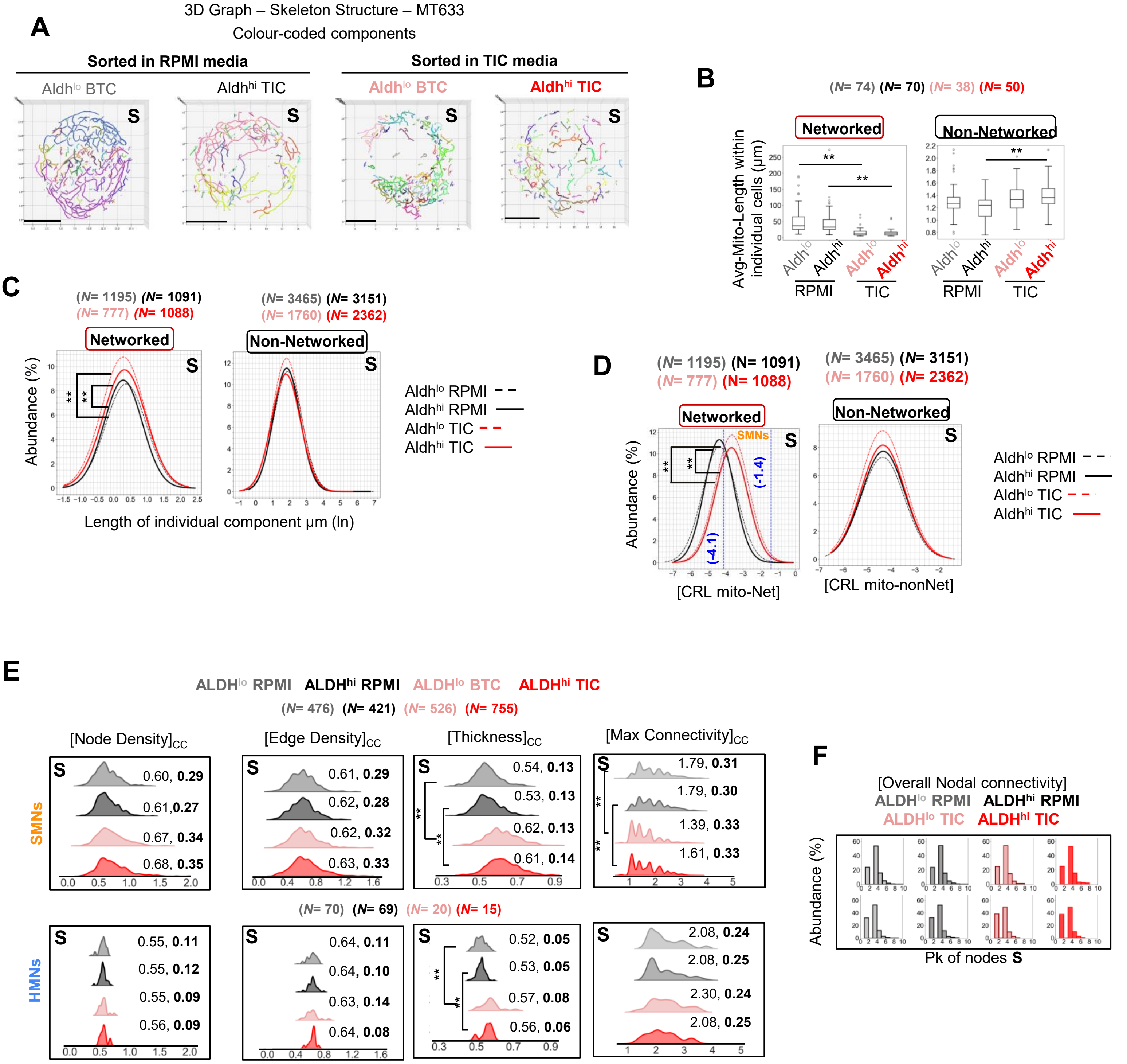
A
3D Graph – Skeleton Structure – MT633
Colour-coded components
Sorted in RPMI media
Sorted in TIC media
Aldhlo BTC
Aldhhi TIC
Aldhlo BTC
Aldhhi TIC
S
B
(N= 74) (N= 70) (N= 38) (N= 50)
Networked
Non-Networked
Avg-Mito-Length within individual cells (µm)
RPMI
TIC
C
(N= 1195) (N= 1091) (N= 777) (N= 1088)
(N= 3465) (N= 3151) (N= 1760) (N= 2362)
Abundance (%)
Length of individual component µm (ln)
Aldhlo RPMI
Aldhhi RPMI
Aldhlo TIC
Aldhhi TIC
D
SMNs
(-1.4)
(-4.1)
[CRL mito-Net]
[CRL mito-nonNet]
E
ALDHlo RPMI ALDHhi RPMI ALDHlo BTC ALDHhi TIC
(N= 476) (N= 421) (N= 526) (N= 755)
[Node Density]CC
[Edge Density]CC
[Thickness]CC
[Max Connectivity]CC
SMNs
0.60, 0.29
0.61,0.27
0.67, 0.34
0.68, 0.35
0.61, 0.29
0.62, 0.28
0.62, 0.32
0.63, 0.33
0.54, 0.13
0.53, 0.13
0.62, 0.13
0.61, 0.14
1.79, 0.31
1.79, 0.30
1.39, 0.33
1.61, 0.33
(N= 70) (N= 69) (N= 20) (N= 15)
HMNs
0.55, 0.11
0.55, 0.12
0.55, 0.09
0.56, 0.09
0.64, 0.11
0.64, 0.10
0.63, 0.14
0.64, 0.08
0.52, 0.05
0.53, 0.05
0.57, 0.08
0.56, 0.06
2.08, 0.24
2.08, 0.25
2.30, 0.24
2.08, 0.25
F
[Overall Nodal connectivity]
ALDHlo RPMI ALDHhi RPMI
ALDHlo TIC ALDHhi TIC
Abundance (%)
Pk of nodes S

**Fig. S4**

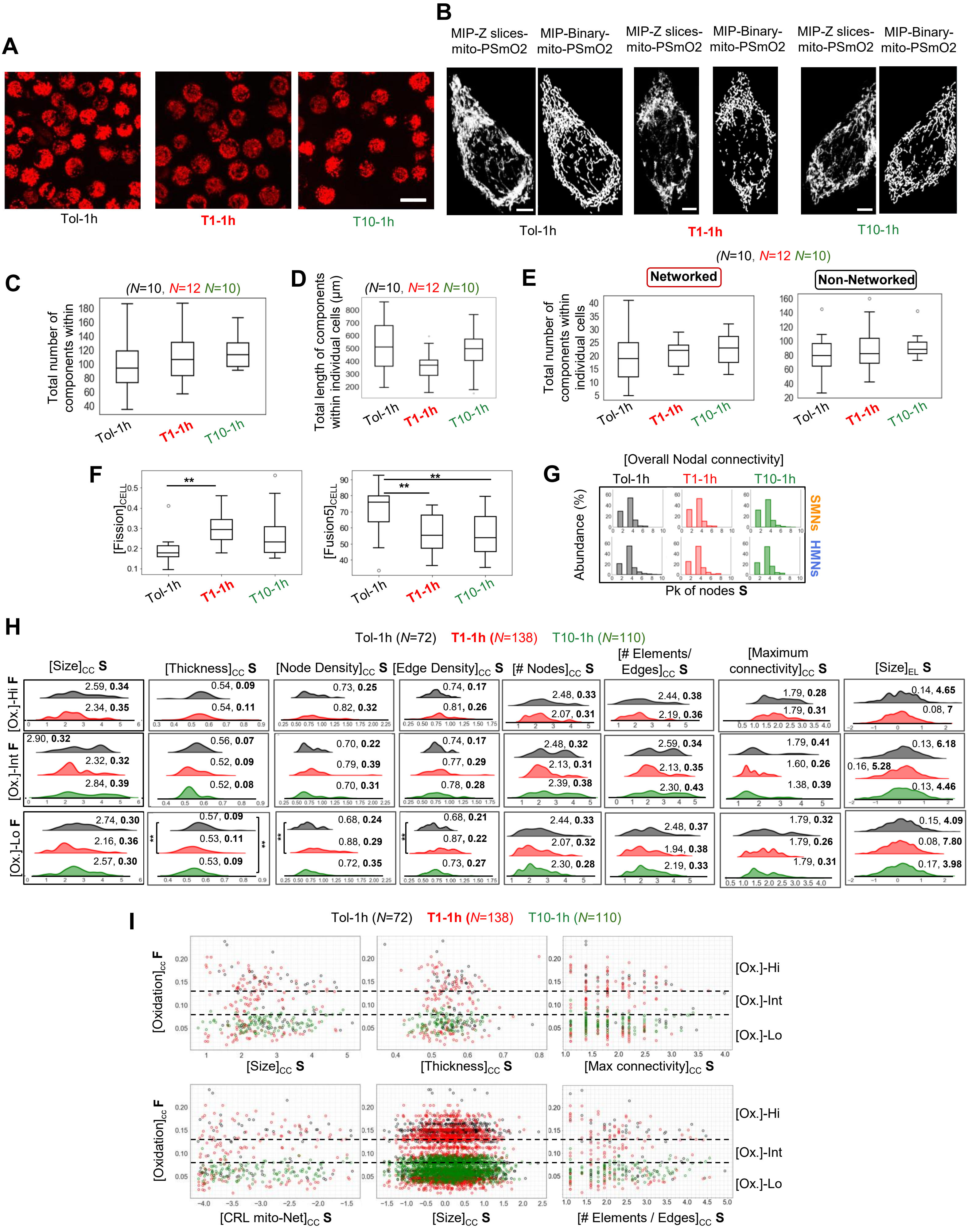
A
Tol-1h
T1-1h
T10-1h
B
MIP-Z slices-mito-PSmO2
MIP-Binary-mito-PSmO2
MIP-Z slices-mito-PSmO2
MIP-Binary-mito-PSmO2
MIP-Z slices-mito-PSmO2
MIP-Binary-mito-PSmO2
Tol-1h
T1-1h
T10-1h
C
(N=10, N=12 N=10)
Total number of components within individual cells
Tol-1h
T1-1h
T10-1h
D
(N=10, N=12 N=10)
Total length of components within individual cells (µm)
Tol-1h
T1-1h
T10-1h
E
(N=10, N=12 N=10)
Networked
Non-Networked
Total number of components within individual cells
Tol-1h
T1-1h
T10-1h
F
[Fission]CELL
[Fusion5]CELL
**
**
**
Tol-1h
T1-1h
T10-1h
G
[Overall Nodal connectivity]
Tol-1h
T1-1h
T10-1h
Abundance (%)
SMNs
HMNs
Pk of nodes S
H
Tol-1h (N=72)
T1-1h (N=138)
T10-1h (N=110)
[Size]CC S
[Thickness]CC S
[Node Density]CC S
[Edge Density]CC S
[# Nodes]CC S
[# Elements/ Edges]CC S
[Maximum connectivity]CC S
[Size]EL S
[Ox.]-Hi F
[Ox.]-Int F
[Ox.]-Lo F
2.59, 0.34
2.34, 0.35
0.54, 0.09
0.54, 0.11
0.73, 0.25
0.82, 0.32
0.74, 0.17
0.81, 0.26
2.48, 0.33
2.07, 0.31
2.44, 0.38
2.19, 0.36
1.79, 0.28
1.79, 0.31
0.14, 4.65
0.08, 7
2.90, 0.32
2.32, 0.32
2.84, 0.39
0.56, 0.07
0.52, 0.09
0.52, 0.08
0.70, 0.22
0.79, 0.39
0.70, 0.31
0.74, 0.17
0.77, 0.29
0.78, 0.28
2.48, 0.32
2.13, 0.31
2.39, 0.38
2.59, 0.34
2.13, 0.35
2.30, 0.43
1.79, 0.41
1.60, 0.26
1.38, 0.39
0.13, 6.18
0.16, 5.28
0.13, 4.46
2.74, 0.30
2.16, 0.36
2.57, 0.30
0.57, 0.09
0.53, 0.11
0.53, 0.09
0.68, 0.24
0.88, 0.29
0.72, 0.35
0.68, 0.21
0.87, 0.22
0.73, 0.27
2.44, 0.33
2.07, 0.32
2.30, 0.28
2.48, 0.37
1.94, 0.38
2.19, 0.33
1.79, 0.32
1.79, 0.26
1.79, 0.31
0.15, 4.09
0.08, 7.80
0.17, 3.98
I
Tol-1h (N=72)
T1-1h (N=138)
T10-1h (N=110)
[Oxidation]cc F
[Size]CC S
[Thickness]CC S
[Max connectivity]CC S
[Ox.]-Hi
[Ox.]-Int
[Ox.]-Lo
[CRL mito-Net]CC S
[Size]CC S
[# Elements / Edges]CC S

Fig.S5

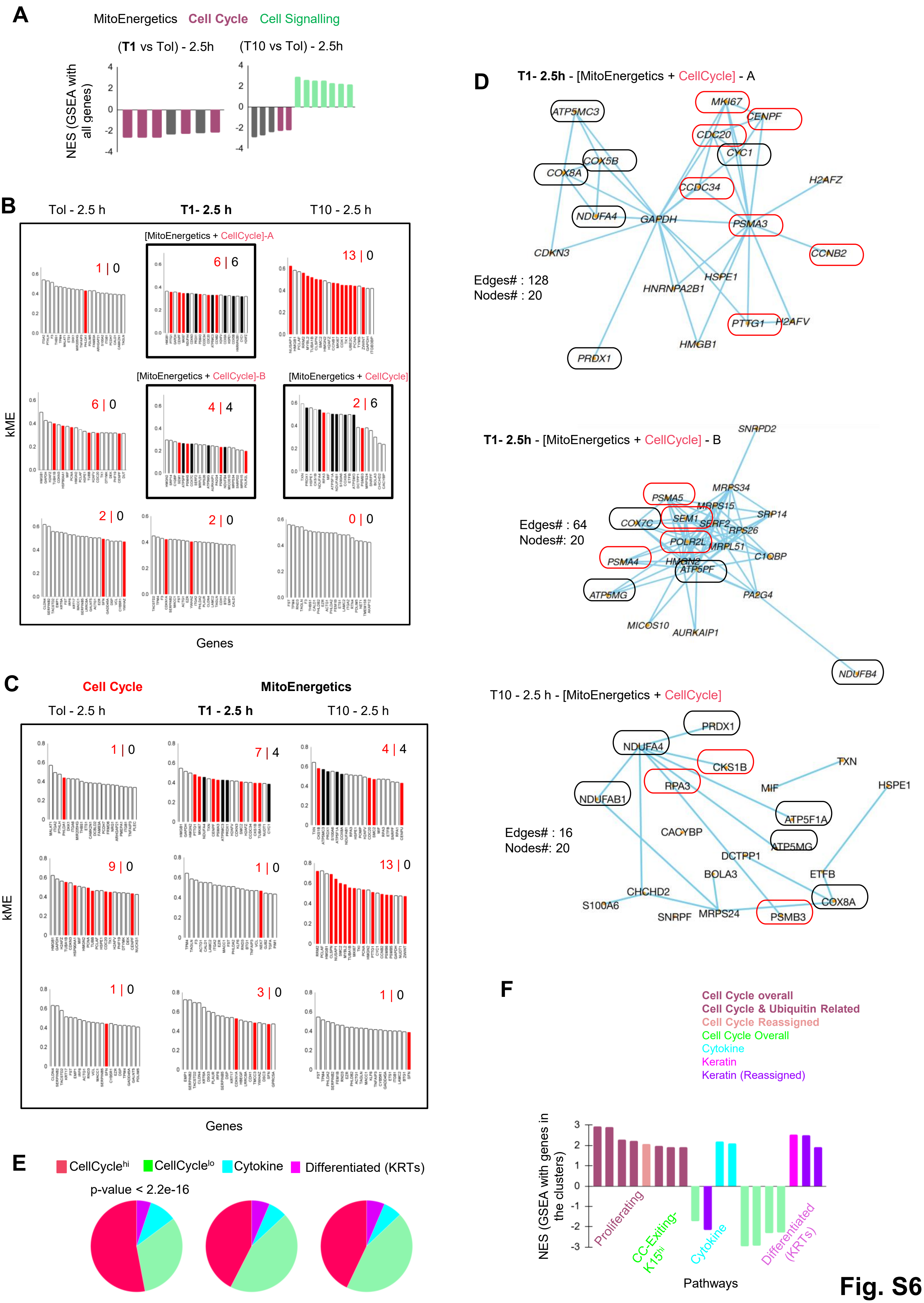
A
MitoEnergetics Cell Cycle Cell Signalling
(T1 vs Tol) - 2.5h
(T10 vs Tol) - 2.5h
NES (GSEA with all genes)
B
Tol - 2.5 h
T1- 2.5 h
T10 - 2.5 h
[MitoEnergetics + CellCycle]-A
[MitoEnergetics + CellCycle]-B
[MitoEnergetics + CellCycle]
1 | 0
6 | 6
13 | 0
6 | 0
4 | 4
2 | 6
2 | 0
2 | 0
0 | 0
kME
Genes
C
Cell Cycle
MitoEnergetics
Tol - 2.5 h
T1 - 2.5 h
T10 - 2.5 h
1 | 0
7 | 4
4 | 4
9 | 0
1 | 0
13 | 0
1 | 0
3 | 0
1 | 0
kME
Genes
D
T1- 2.5h - [MitoEnergetics + CellCycle] - A
Edges# : 128
Nodes# : 20
ATP5MC3
MKI67
CENPF
CDC20
CYC1
COX5B
COX8A
CCDC34
H2AFZ
NDUFA4
GAPDH
PSMA3
CDKN3
CCNB2
HSPE1
HNRNPA2B1
PTTG1
H2AFV
HMGB1
PRDX1
T1- 2.5h - [MitoEnergetics + CellCycle] - B
Edges# : 64
Nodes#: 20
SNRPD2
MRPS34
PSMA5
MRPS15
COX7C
SEM1
SERF2
SRP14
RPS26
POLR2L
MRPL51
PSMA4
HMGN2
C1QBP
ATP5PF
ATP5MG
PA2G4
MICOS10
AURKAIP1
NDUFB4
T10 - 2.5 h - [MitoEnergetics + CellCycle]
Edges# : 16
Nodes#: 20
PRDX1
NDUFA4
CKS1B
TXN
RPA3
MIF
HSPE1
NDUFAB1
ATP5F1A
CACYBP
ATP5MG
DCTPP1
BOLA3
ETFB
CHCHD2
COX8A
S100A6
SNRPF
MRPS24
PSMB3
E
CellCycle^hi CellCycle^lo Cytokine Differentiated (KRTs)
p-value < 2.2e-16
F
Cell Cycle overall
Cell Cycle & Ubiquitin Related
Cell Cycle Reassigned
Cell Cycle Overall
Cytokine
Keratin
Keratin (Reassigned)
NES (GSEA with genes in the clusters)
Proliferating
CC-Exiting-K15hi
Cytokine
Differentiated (KRTs)
Pathways

Fig. S6

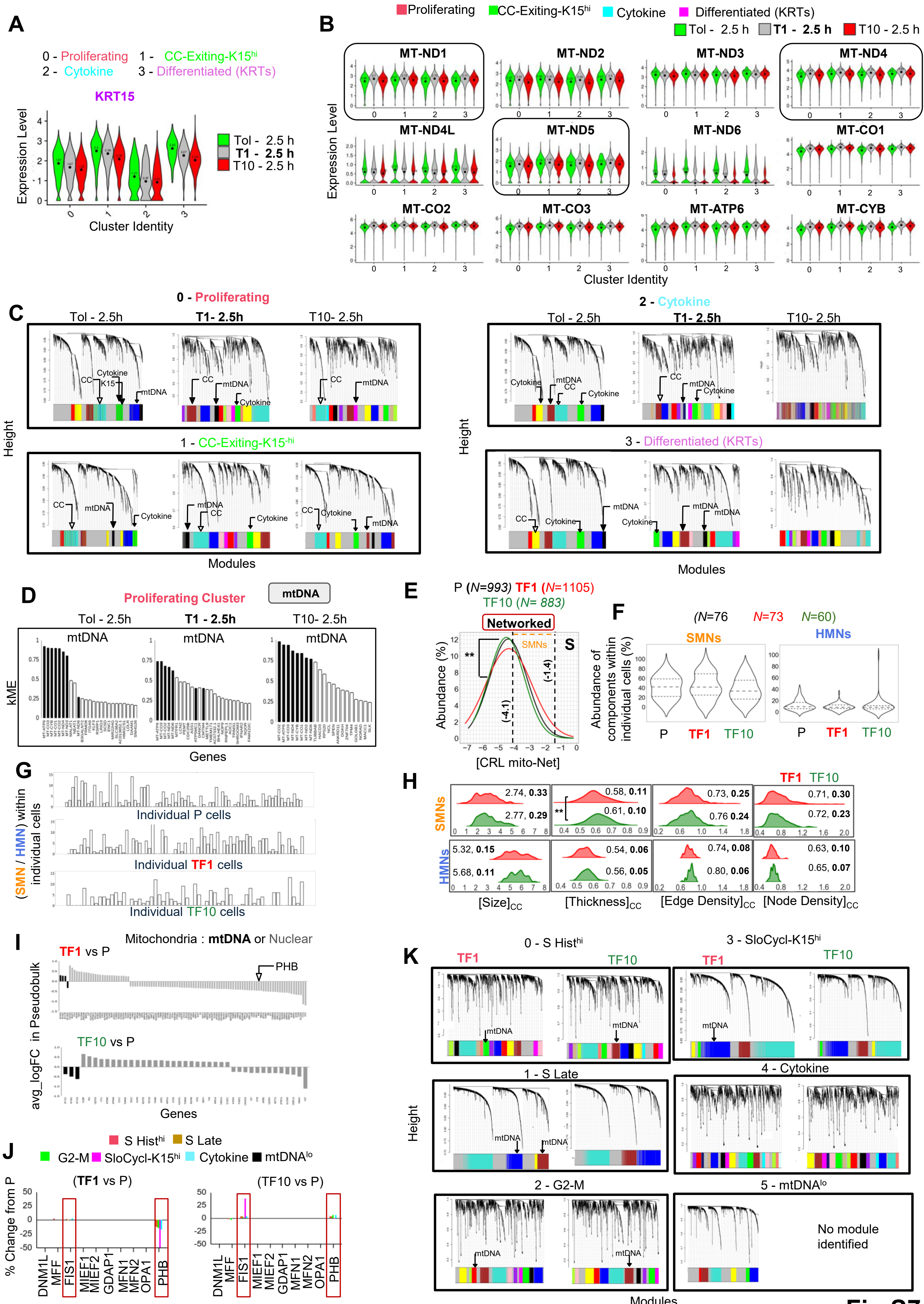
A
0 - Proliferating 1 - CC-Exiting-K15hi
2 - Cytokine 3 - Differentiated (KRTs)
KRT15
Expression Level
Tol - 2.5 h
T1 - 2.5 h
T10 - 2.5 h
Cluster Identity
B
Proliferating CC-Exiting-K15hi Cytokine Differentiated (KRTs)
Tol - 2.5 h T1 - 2.5 h T10 - 2.5 h
MT-ND1
MT-ND2
MT-ND3
MT-ND4
MT-ND4L
MT-ND5
MT-ND6
MT-CO1
MT-CO2
MT-CO3
MT-ATP6
MT-CYB
Expression Level
Cluster Identity
C
0 - Proliferating
Tol - 2.5h T1- 2.5h T10- 2.5h
2 - Cytokine
1 - CC-Exiting-K15-hi
3 - Differentiated (KRTs)
CC
K15
Cytokine
mtDNA
Height
Modules
D
Proliferating Cluster
mtDNA
Tol - 2.5h T1 - 2.5h T10- 2.5h
kME
Genes
E
P (N=993) TF1 (N=1105)
TF10 (N= 883)
Networked
SMNs
S
**
(-4.1)
(-1.4)
Abundance (%)
[CRL mito-Net]
F
(N=76 N=73 N=60)
SMNs
HMNs
Abundance of components within individual cells (%)
P TF1 TF10
G
(SMN / HMN) within individual cells
Individual P cells
Individual TF1 cells
Individual TF10 cells
H
TF1 TF10
SMNs
HMNs
2.74, 0.33
2.77, 0.29
0.58, 0.11
0.61, 0.10
0.73, 0.25
0.76 0.24
0.71, 0.30
0.72, 0.23
5.32, 0.15
5.68, 0.11
0.54, 0.06
0.56, 0.05
0.74, 0.08
0.80, 0.06
0.63, 0.10
0.65, 0.07
[Size]CC
[Thickness]CC
[Edge Density]CC
[Node Density]CC
I
Mitochondria : mtDNA or Nuclear
TF1 vs P
PHB
TF10 vs P
avg_logFC in Pseudobulk
Genes
J
S Histhi S Late
G2-M SloCycl-K15hi Cytokine mtDNAlo
(TF1 vs P)
(TF10 vs P)
% Change from P
DNM1L
MFF
FIS1
MIEF1
MIEF2
GDAP1
MFN1
MFN2
OPA1
PHB
Fission Fusion Genes
K
0 - S Histhi
3 - SloCycl-K15hi
TF1
TF10
mtDNA
1 - S Late
4 - Cytokine
2 - G2-M
5 - mtDNAlo
No module identified
Height
Modules

Fig.S7

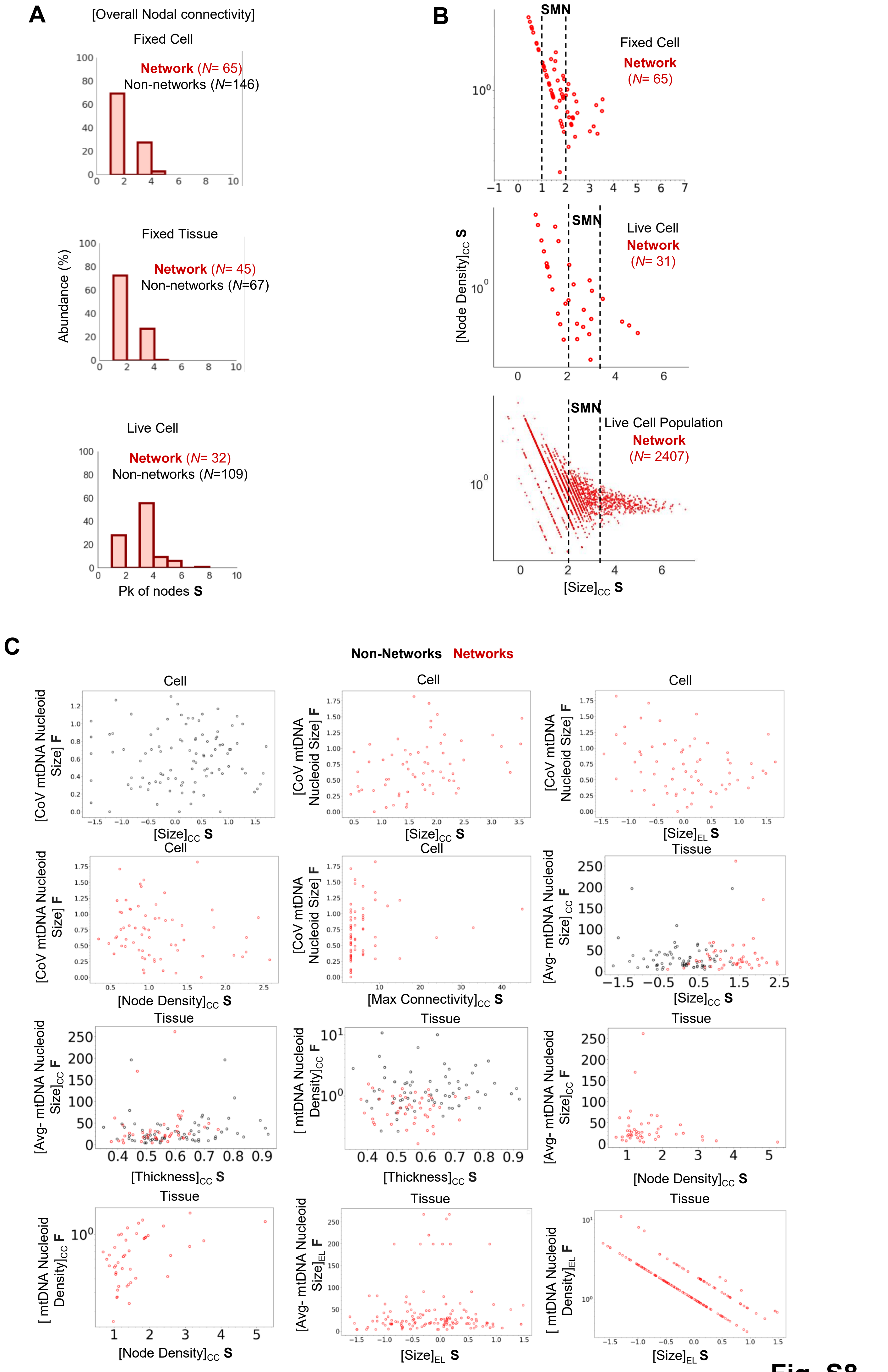
A
[Overall Nodal connectivity]
Fixed Cell
Network (N= 65)
Non-networks (N=146)
Fixed Tissue
Network (N= 45)
Non-networks (N=67)
Live Cell
Network (N= 32)
Non-networks (N=109)
Abundance (%)
Pk of nodes S
B
SMN
Fixed Cell
Network (N= 65)
Live Cell
Network (N= 31)
Live Cell Population
Network (N= 2407)
[Node Density]CC S
[Size]CC S
C
Non-Networks Networks
Cell
Tissue
[CoV mtDNA Nucleoid Size] F
[Avg- mtDNA Nucleoid Size]CC F
[mtDNA Nucleoid Density]CC F
[Avg- mtDNA Nucleoid Size]EL F
[mtDNA Nucleoid Density]EL F
[Size]CC S
[Size]EL S
[Node Density]CC S
[Max Connectivity]CC S
[Thickness]CC S

Fig. S8

**A**

| line_id | point_id | x | y | z | x_pixel | y_pixel | z_pixel |
|---|---|---|---|---|---|---|---|
| 0 | 0 | 8.97867 | 4.85333 | 2.33333 | 86 | 47 | 5 |
| 0 | 1 | 8.85002 | 4.95192 | 2.41975 | 85 | 48 | 5 |
| 0 | 2 | 8.73600 | 4.99200 | 2.50000 | 84 | 48 | 5 |
| 1 | 0 | 17.42000 | 6.05800 | 2.37500 | 168 | 58 | 5 |
| 1 | 1 | 17.27913 | 5.89616 | 2.43519 | 166 | 57 | 5 |

**B**

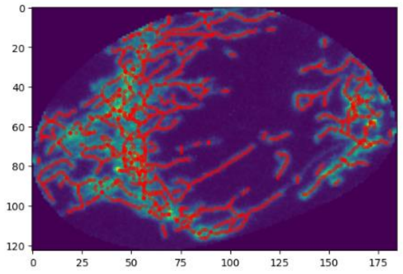

**C**

| x_pixel | y_pixel | z_pixel | pixel_intensity_channel_1 | pixel_intensity_channel_2 | cc_pixel_intensity_ratio |
|---|---|---|---|---|---|
| 105 | 55 | 17 | 256.5 | 12.75 | 0.05407279 |
| 106 | 54 | 17 | 243.5 | 35 | 0.05407279 |
| 107 | 54 | 18 | 374 | 18 | 0.05407279 |
| 108 | 54 | 18 | 483.5 | 34.25 | 0.05407279 |
| 109 | 55 | 18 | 463.5 | 17 | 0.05407279 |
| 110 | 56 | 18 | 342.75 | 0 | 0.05407279 |

**D**

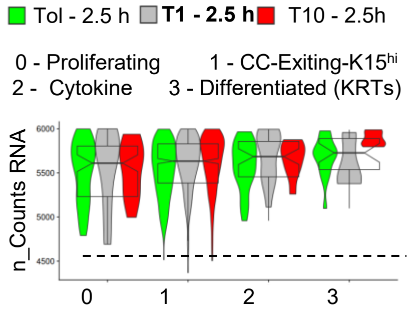


**E**

| Dataset | nCount_RNA | KNN | Max shared |
|---|---|---|---|
| Acute -2.5h: Cell Clusters | >=2000 | 20 | 10 |

**F**

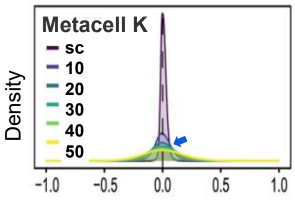


**G**

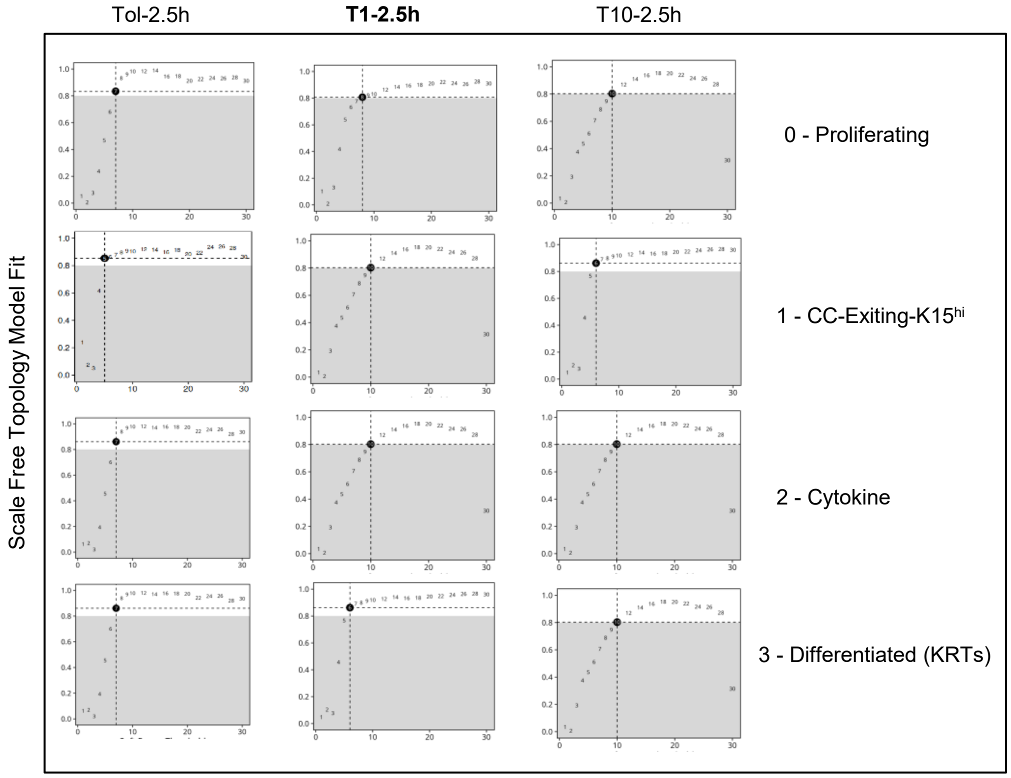


**Fig. S9**

**Supplementary Fig. S1:**

**(A).** Scatter plot of total length of mitochondrial components within individual cells calculated using mito-SinComp analysis and directly from Mitograph**.**

**(B).** Scatter plot of $[Fission]_{CELL}$ versus $[Fusion5]_{CELL}$ metrics from mito-SinCe analysis of individual cells stained with MTG in a combined population with varying levels of Drp-1 knockdown {re-analysed from Ref. 23}; range for INT-cells and HF-cells are demarcated by coloured boxes.

(**C**)**.** Box plot of abundance of mitochondrial components within individual cells in categories mentioned.

(**D**)**.** Box plot of average mitochondrial length within individual cells in categories mentioned.

**(E).** Box plot of the total number of mitochondrial components within individual cells in categories mentioned.

**(F).** Box plot of the total length of mitochondrial components within individual cells ($[Sum\ L_{COMP}]_{CELL}$) in categories mentioned.

(**G**)**.** Bivariate scatter plot of $[Oxidation]_{CELL}$ values obtained using automated mito-SinComp structure-function script and that manually from Zen-blue/black software for averaged values at the single-cell level.

(**H**)**.** Box plot of the $[Oxidation]_{CELL}$, as calculated using the mito-SinComp script (left) and manually using image analysis software (right).

(**I**)**.** Ridgeline plots comparing the distribution of $[Oxidation]_{CC}$ between Networked and non-Networked mitochondrial components in the individual experimental groups from **Fig 2E**; Median and Coefficient of Variation are indicated for each feature latter in bold.

**signifies Holmes-Bonferroni corrected p-value < 0.05; N = sample size.

**Supplementary Fig. S2:**

**(A).** S vs F bivariate scatter plots of INH-$[Oxidation]_{CC}$ and individual structural features with their frequency distributions from the pooled cell groups of **I**; dotted lines depict distributions and coloured dots depict cell groups.

**(B).** Magnified view of the extracted networks and non-networks from the cell in (**Fig 2G**), with corresponding micrographs along with S and F feature descriptions (red denotes the respective median values).

**Supplementary Fig. S3:**

(**A**)**.** Histogram plots representing overall nodal connectivity of SMNs and HMNs.

(**B**)**.** Workflow for detailed stepwise process for hdWGCNA. The hdWGCNA workflow allows network-based transcriptomic analysis in both pseudobulk and single-cell cluster contexts. The stepwise process begins with setting up hdWCGNA object, then constructing metacells using K-nearest neighbors (KNN), followed by computing Pearson's correlations, and transforming them into adjacency matrices with soft-thresholding. Thereafter, networks are built using Topological Overlap Measures (TOM), and transcriptomic modules are identified via hierarchical clustering. Finally, module eigengenes (ME) and connectivity (kME) values are calculated to identify hub genes.

**(C).** Dendrograms depicting hierarchical clustering of adjacency-based dissimilarity to identify color-coded co-expression modules, where grey module indicates no significant co-expression.

**(D).** kME plot of KRT15 transcriptomic interaction modules identified using hdWGCNA, of the pairwise pseudo-bulk analyses gene set of the pooled scRNA-seq data set of HaCAT cells with weak or strong knockdown of Drp1 along with its non-targeting control (Original data from Ref. 23).

(**E**)**.** Hub-Gene Network plot for the KRT15 transcriptomic interaction module of **D**; direct KRT15 (purple outline) transcriptomic connections are highlighted (dashed outlines).

**(F).** Table depicts edge distances of the direct transcriptomic connections of KRT15 outlined in **E**; candidates in blue are common with cluster analyses in **Fig 3E**.

(**G**)**.** Violin plots depicting gene expression of KRT15 and its direct transcriptomic connections across scRNA-seq derived clusters in the DRP1-kd (W), DRP1-kd (S) and NT (control).

**Supplementary Fig. S4:**

**(A).** Representative 3D structure Graphs of Mitotracker 633 stained single A2780-CP cells from the $Aldh^{lo}$ BTCs and $Aldh^{hi}$ TICs in RPMI (growth media) or TIC (stem cell media) groups.

(**B**)**.** Box plot of average mitochondrial length within individual cells in the categories mentioned.

**(C).** Histogram distribution of lengths of individual networked and non-networked mitochondrial components in categories mentioned.

**(D).** Histogram distribution of [CRL mito-Net] and [CRL mito-nonNet] feature in the color-coded groups mentioned, with demarcation of thresholds (dashed line) for SMNs and HMNs (as in **Fig. 3A**).

**(E).** Ridgeline plots comparing the distribution of the named structural features of SMNs and HMNs between the color-coded groups; Median and Coefficient of Variation are indicated for each feature, latter in bold.
**(F).** Histogram plots representing overall nodal connectivity of SMNs and HMNs in color coded groups.

**signifies Holmes-Bonferroni corrected p-value < 0.05; Scale bar: 5 μm; N = sample size.

**Supplementary Fig. S5:**
**(A).** Maximum Intensity Projections of Z-slices of TMRE-stained HaCaT cells after Tol-1h, T1-1h, T10-1h treatment. Scale bar: 20 μm
**(B).** Labelled views of HaCaT cells stably expressing mito-PSmO2 from the groups mentioned. Scale bar: 5 μm.
(**C**)**.** Box plot of the total number of mitochondrial components (including networks and non-networks) within individual cells from the groups mentioned.
(**D**)**.** Box plot of the total length of mitochondrial components (including networks and non-networks) within individual cells from the groups mentioned.
**(E).** Box plot of the total number of mitochondrial components within individual cells in categories mentioned.
**(F).** Box plot of $[\text{Fission}]_{\text{CELL}}$ and $[\text{Fusion5}]_{\text{CELL}}$ metrics of individual cells from mito-SinCe analyses of labelled groups.
(**G**)**.** Histogram plots representing overall nodal connectivity of SMNs and HMNs in color coded groups.
(**H**)**.** Ridgeline plots compare the distribution of the named structural features of SMNs between the color-coded groups; Median and Coefficient of Variation are indicated for each feature.
**(I).** Various S-F bivariate scatter plots of SMNs in the color-coded groups. The horizontal dashed lines indicate the demarcation of the S and F features, respectively.

**signifies Holmes-Bonferroni corrected p-value < 0.05. N = sample size.

**Supplementary Fig. S6:**
**(A).** Normalized Enrichment Scores (NES) from Gene Set Enrichment Analysis (GSEA) of all genes from pseudo-bulk analyses described in **Fig. 5A**; enriched pathways are color coded.

(**B**). kME plot of modules identified by hdWGCNA using all genes from T1 vs Tol pairwise pseudobulk analysis; [MitoEnergetics + CellCycle] modules are boxed.
(**C**). kME plot of modules identified by hdWGCNA using all genes from T10 vs Tol pairwise pseudobulk analysis.
(**D**). Hub-Gene Network plots of modules identified in **B** (top20); mitochondrial and cell genes are outlined.
(**E**). Pie chart displaying the percentage distribution of scRNA-seq derived cell clusters in the cell populations identified in **Fig. 5C**.
(**F**). Normalized enrichment scores of functional pathways identified by GSEA for cluster markers. Functional annotations based on leading-edge genes are listed on the right, with color-coded clusters arranged as in **Fig. 5C**

**Supplementary Fig. S7:**
**(A).** Violin plots showing expression levels of the stem cell gene KRT15 in Tol-2.5h, T1-2.5h and T10-2.5h cell populations across scRNA-seq derived cell clusters.
**(B).** Violin plots showing the expression levels of mtDNA genes in Tol-2.5h, T1-2.5h and T10-2.5h cell populations across scRNA-seq derived cell clusters.
(**C**). Dendrograms depicting distinct co-expression modules in various scRNA-seq derived cell clusters (0-3) for Tol-2.5h, T1-2.5h and T10-2.5h populations; arrows indicate GSEA based functional annotations.
(**D**). kME plots showing transcriptomic interaction modules obtained using hdWGCNA in Tol-2.5h, T1-2.5h and T10-2.5h cell populations across scRNA-seq derived cell clusters.
**(E).** Histogram distribution of [CRL mito-Net] feature in the Parental (P), TCDD-1nM and TCDD-10nM transformed (TF1 and TF10) HaCaT cells, with demarcation of thresholds (dashed line) for SMNs (as in **Fig. 3A**).
**(F).** Violin plot showing abundance of mitochondrial networks per cell in SMN and HMN categories in color coded experimental groups, with median (dashed line) and the quartile ranges (dotted lines) shown
**(G).** Bar plot showing (SMN / HMN) ratio within individual cells among color-coded groups.
**(H).** Ridgeline plots comparing the distribution of the named structural features between SMNs and HMNs from TF1 and TF10 groups; Median and Coefficient of Variation are indicated for each parameter, latter in bold.
**(I).** Pairwise pseudo-bulk analyses on scRNA-seq data from TF1 and TF10 populations in comparison to Parental (P).

**(J).** Bar plots showing the percentage change in median expression of Fission-Fusion genes in experimental groups (TF1 and TF10) relative to the control (P) across distinct cell clusters. FIS1 and PHB are outlined.
**(K).** Dendrograms depicting distinct co-expression modules in various scRNA-seq derived cell clusters (0-5) for TF1 and TF10 populations; arrows indicate GSEA based functional annotations.

**Supplementary Fig. S8:**
**(A).** Comparison of histogram plots representing overall nodal connectivity of the groups mentioned in the representative fixed cell (**Fig. 6**) and representative live cells (**Fig. 2**).
**(B).** Bivariate S-F scatter plot of $[\text{Node density}]_{CC}$ with $[\text{Size}]_{CC}$ of the groups mentioned, with SMN demarcations shown based on mean and median of $[\text{Size}]_{CC}$ in live cell populations.
**(C)** Various S-F bivariate scatter plots from mito-SinComp analyses on fixed cells and tissues.

**Supplementary Fig. S9:**
**(A).** Tabular representation of conversion of MitoGraph output to pixel coordinates.
**(B).** Mapping of calculated pixel coordinates over structure. Image of a representative cell showing calculated pixel coordinates (red) from MitoGraph overlayed on the MIP of the structural channel for individual mitochondrial components (green).
**(C).** Tabular representation of individual pixel coordinates and their corresponding pixel intensity value to obtain pixel intensity ratio reflecting redox function for a representative individual non-network (cc).
**(D).** Violin plots display the distribution of high-quality cell counts(n_Counts RNA) for Tol-2.5h, T1-2.5h and T10-2.5h across scRNA-seq derived clusters; dashed lines represent the selected threshold.
**(E).** Table with parameters for n_Counts RNA, KNN and maximum shared (max shared) considered for the designated Tol-2.5h, T1-2.5h and T10-2.5h cell populations.
**(F).** Density plot illustrating the distribution of pairwise Pearson correlations between genes from the single-cell expression matrix and metacell expression matrices with varying values of KNN parameter K; blue arrow indicates the selected KNN value.
**(G).** Optimal soft-thresholding power ($\beta$) for Tol-2.5h, T1-2.5h and T10-2.5h pooled data for the scale-free topology model fit index for hdWCGNA objects. Numbers mentioned within solid black circles indicate the selected $\beta$ value.

Supplemental Table 1

Click here to access/download
**Supplemental File Sets**
Supp.Table 1-09-06-25-Submit.docx

Supplemental Table 2

Click here to access/download
**Supplemental File Sets**
Supp.Table 2-09-06-25-Submit.docx
